# Measuring mRNA translation in neuronal processes and somata by tRNA-FRET

**DOI:** 10.1101/646216

**Authors:** Bella Koltun, Sivan Ironi, Noga Gershoni-Emek, Iliana Barrera, Mohammad Hleihil, Siddharth Nanguneri, Ranjan Sasmal, Sarit S. Agasti, Deepak Nair, Kobi Rosenblum

## Abstract

In neurons, the specific spatial and temporal localization of protein synthesis is of great importance for function and survival. In this work, we visualized tRNA and protein synthesis events in fixed and live mouse primary cortical culture using fluorescently-labeled tRNAs. We were able to characterize the distribution and movement of tRNAs in different neuronal sub-compartments and to study their association with the ribosome. We found that tRNA motion in neural processes is lower than in somata and corresponds to patterns of slow transport mechanisms, and that larger tRNA puncta co-localize with translational machinery components and are likely the functional fraction. Furthermore, chemical induction of LTP in culture revealed GluR-dependent biphasic up-regulation of mRNA translation with a similar effect in dendrites and somata. Importantly, measurement of protein synthesis in neurons with high resolutions offers new insights into neuronal function in health and disease states.

## Introduction

Proteostatic processes, including protein synthesis and/or degradation and the mechanisms regulating them, are key to the cellular ability to respond to environmental changes. In neurons, the spatiotemporal localization of mRNA translation is of paramount importance due to their unique morphology. In neuronal processes, which often traverse distances several orders of magnitude larger than the cell body, the regulation of local translation is fundamental for maintaining their distinct functionalities, including signaling and synaptic plasticity (1, 11, 94, 99). Moreover, *de novo* protein synthesis is required for the formation of long-term memory (consolidation of labile, short term memory to more stable, long term memory) as well as synaptic plasticity (56–58). At the same time, dysregulation of these mechanisms underlies various neurodevelopmental and neurodegenerative pathologies. Both the initiation and elongation phases of mRNA translation and the respective translation factors that mediate them have been suggested as innovative targets for memory enhancement in health and disease (56, 59–63, 111).

Local protein synthesis has been at the forefront of neuroscience for many years, originating in the discovery of polyribosomes at the base of dendritic spines (98). Over the years, a combination of methodologies, including *in situ* hybridization, deep sequencing, and microarrays have shown that neurites possess a large transcriptome that is subject to alterations as a result of development, disease, and environment (81, 95, 109). The translational efficiency of mRNAs *in vivo* can be studied using ribosome profiling, which has also revealed valuable information regarding translation regulation. However, these methods are limited temporally and require the averaging of many cells, which masks individual cellular contributions (83, 87, 89, 92, 100). In recent years, together with the recognition of the importance of a single cell within a population, imaging techniques have taken central stage. Several methodologies have been developed in order to probe local protein synthesis. Average synthesis rates were acquired from single-cell imaging (84, 91, 100), subcellular sites of translation were recognized by co-localizing mRNAs and ribosomes (9, 40, 90), and the initial translation event of a single mRNA was also observed (36). More recently, Cre-dependent conditional expression of non-canonical amino acids was used to label nascent proteomes *in vivo* in a cell-type-specific manner (54).

mRNA translation is a cyclical process consisting of initiation, elongation, and termination. Initiation is commenced by the binding of the initiator methionyl-tRNA (Met-tRNA) to the 40s ribosomal subunit in a ternary complex together with the GTP-bound eukaryotic initiation factor eIF2 to create the pre-initiation complex, and culminates in the recognition of the mRNA start codon by the functional 80s subunit (11, 97). During elongation, the Met-tRNA is base-paired with the AUG start codon at the P site, while the next codon awaits at the A-site. The recognition of the codon by the tRNA triggers GTP hydrolysis, while peptide formation occurs in the peptidyl site of the ribosome. Thus, the polypeptide is transferred from the P site to the A site, and the nascent protein is extended by one amino acid (82).

tRNAs are the most abundant small non-coding RNA species in the cell, making up about 10% of all cellular RNAs. They have a canonical role as an adaptor molecule during protein synthesis, in addition to various non-translational roles (46). tRNA is a 76-90 nucleotide molecule which is transcribed from hundreds of different genes (497 known in humans) scattered throughout the genome. These genes give rise to up to 46 different tRNA isoacceptors in one human cell, capable of base-pairing with the 61 known sense codons that comprise the genetic code (due to the wobble base in the first position of the anticodon). The three-dimensional L-shape of the tRNA is highly conserved, consisting of two helices: the acceptor stem is stacked on top of the TψC loop, and the anticodon stem is stacked on top of the D-loop (80).

Different cell types within an organism differ in their tRNA expression profiles. Studies have shown a correlation between the abundance of specific tRNAs and the enrichment of their corresponding codon in the cell transcriptome (88), as well as tissue-specific tRNA expression profiles (17). This suggests a role for tRNA in cellular identity and function, however, little is known about their formation, localization, and function in neurons. All of these can putatively regulate tRNA involvement in mRNA translation, and thus regulate protein synthesis.

Here, we present a novel method for inspecting both tRNA dynamics and mRNA translation by fluorescently-tagged tRNAs in sub-neuronal compartments. As tRNAs directly link the mRNA to the amino acid within the ribosome, they are an attractive candidate for imaging mRNA translation *in vitro*. Bulk yeast tRNA, fluorescently-labeled on its D-loop, was introduced into cortical primary cultures and used to measure tRNA distribution and characteristics. The nucleic acid sequence and three-dimensional structure of tRNA are highly evolutionarily conserved, as is the codon-anticodon system. Fluorescently-labeled yeast tRNA is functional and capable of mediating mRNA translation in higher eukaryotes (8, 53). Using super-resolution microscopy, we analyzed tRNA localization in different neuronal compartments and showed that a fraction of fluorescently-labeled tRNA associates with assembled ribosomes in dendritic spines.

We used two tRNA populations, which were labeled with Cy3 and Cy5 dyes, and thus enabled us to measure Förster Resonance Energy Transfer (FRET). FRET can only occur when two differently-labeled tRNAs are located at the A and P sites of an active ribosome, enabling us to reliably monitor active mRNA translation in neurons (8). We found that the level of mRNA translation is similar in neuronal somata and processes and is negatively correlated with the developmental age of neuronal cultures. Moreover, following chemical LTP induction, we detected two waves of mRNA translation up-regulation both in neuronal somata and processes. Altogether, our data show that the tRNA-FRET method can serve as a reliable reporter for mRNA translation in live neurons, and can be used to visualize translation with precise spatiotemporal resolution.

## Materials and methods

### Animals

For primary culture, C57BL/6 WT pregnant dams 2-3 months of age were obtained weekly from local vendors (Envigo RMS, Jerusalem, Israel). *Arc:dVenus* mice (with a C57BL/6 background), in which an extra chromosome was added with the dVenus gene reporter under the regulation of the Arc promoter, were generously provided by S.A. Kushner. dVenus-positive mice and wild-type (WT) littermates were derived by crossing heterozygous and wild-type mice. Transgenic *Arc:dVenus* mice were identified with the following primer sequences, which yielded a 560bp band:

F: 5’-CTGACCCTGAAGCTGATCT-3’

R: 5’-AGGGTACAGCTCGTCCAT-3’

For electrophysiology experiments, 2-3 months old C57BL/6 WT mice were used. Mice were maintained on a 12 h light/dark cycle with unlimited food and water availability. All experiments were approved by the University of Haifa Animal Care and Use Committee and were in accordance with the National Institutes of Health guidelines for the ethical treatment of animals (ethics #485/17, #566/18, #553/18).

### Primary cortical culture

P1 neonatal C57BL/6 and *Arc:dVenus* mouse cortices were used to prepare primary cultures as previously described in (61). Primary cultures were seeded in non-pyrogenic polystyrene cell culture plates (Corning, Corning, NY). For immunocytochemistry, 18 mm #1 coverslips (ThermoFisher Scientific, Waltham, MA) in 12-well plates were coated with 0.01% polyornithine (#P3655, Sigma-Aldrich, Rehovot, Israel) together with 2µg/mL laminin (#L2020, Sigma-Aldrich) and incubated at 37°C for 4 h prior to seeding, and 87,000 cells were seeded in 1 mL. For western blot analysis, 6-well plates were coated with 0.2% polyethynelimine and incubated overnight at 4°C prior to seeding, and 300,000 cells were seeded in 2 mL. Medium was replaced with B-27 (#17504044, ThermoFisher Scientific)/MEM (diluted according to manufacturer’s instructions) for increased neuronal survival after 6-8 DIV.

For axon imaging, 300,000 cells were seeded in silicon microfluidic chambers, a gift from E. Perlson, Tel Aviv University (Figure 5 – figure supplement 1A). Cell bodies were seeded in a proximal chamber, and axons grew through grooves 300 µm in length towards the distal side. After axon crossing to the distal chamber (approximately 5-7 days), cells were transfected with tRNA and imaged two to seven days later.

### Introduction of labeled tRNA

Bulk tRNA was labeled with either Cy3 or Cy5 fluorescent dye (Anima Biotech, Bernardsville, NJ) (8, 53). Cells were transfected with 100 ng labeled tRNA for 87,000 cells, using 1µl of PolyJet transfection reagent (#SL100688, SignaGen Laboratories, Rockville, MD) according to manufacturer’s instructions. All cultures were transfected 48 hours prior to fixation/live imaging and imaged at 12-15 DIV unless stated otherwise.

### Electrophysiology

2-3 months old male C57BL/6 WT mice were anesthetized with isoflurane. Following decapitation, brains were quickly removed and immersed in cooled (1-3°C), oxygenated (95% O_2,_ 5% CO_2_) sucrose-based cutting solution containing the following (in mM): 110 sucrose, 60 NaCl, 3 KCl, 1.25 NaH_2_PO_4_, 28 NaHCO_3_, 0.5 CaCl_2_, 7 MgCl_2_, 5 D-glucose, and 0.6 ascorbate (all chemicals obtained from Sigma-Aldrich). The brain tissue containing the hippocampus was dissected, glued to a small stage using Cyanoacrylates-based glue and immersed in the cutting solution. Coronal slices 300 µm thick were made using a vibratome (model 7000, Campden Instruments, Loughborough, UK) in cooled, oxygenated cutting solution. The slices were allowed to recover for 30 min at 37°C in artificial CSF (ACSF) containing the following (in mM): 125 NaCl, 2.5 KCl, 1.25 NaH_2_PO_4_, 25 NaHCO_3_, 25 D-glucose, 2 CaCl_2_, and 1 MgCl_2_ (Sigma-Aldrich), followed by additional recovery for at least 30 min in ACSF at room temperature until electrophysiological recording. After the recovery period, slices were placed in the recording chamber and maintained at 34°C with continuous perfusion of carbonated ACSF (95% O_2_ 5% CO_2_).

Recordings were amplified by Axopatch 200B amplifier and digitized with Digidata 1440 (Molecular Devices, San Jose, CA). The recording electrode was pulled from a borosilicate glass pipette (3–5 MΩ; #B150-110-10HP, Sutter Instruments, Novato, CA) using an electrode puller (#P-1000; Sutter Instruments) and filled with a K-gluconate-based internal solution containing the following (in mM): 135 K-gluconate, 4 KCl, 10 HEPES, 10 phosphocreatine, 4 Mg-ATP, 0.3 Na_3_GTP, (all reagents obtained from Sigma Aldrich), osmolarity 290 mOsm, pH 7.3. tRNA was added to the pipette solution (100 ng) and whole cell current clamp recordings were made from the soma of CA1 pyramidal cells.

Resting membrane potential was estimated by measuring the voltage in the absence of any current injection after 5 min of achieving whole cell configuration. The dependence of firing rate on the injected current was obtained by injection of current steps (of 500ms duration from 50 to 400 pA in 50 pA increments). Input resistance and membrane time constant were calculated from the voltage response to a hyperpolarizing current pulse (150 pA). To measure action potential threshold, current pulses (5pA, 10ms) were injected from 70mV holding potential, a curve of dV/dt was created for the first action potential (AP) that appeared on the 5 ms time point, and the 30 V/s point in the rising slope of the AP was considered as threshold. To standardize adaptation recordings, 1 sec depolarizing current steps were injected to the cell body to determine the current intensity needed to generate a single action potential (I-thr). Action potential adaptation ratio was calculated from 1 sec current step with stimulus intensity equal to I-thr X2. The firing frequency at each inter-spike interval (ISI) along the train was normalized to the initial frequency at the train onset. Liquid junction potential was not corrected. Recordings were low-pass filtered at 10 kHz and sampled at 20 kHz. Pipette capacitance and series resistance were compensated and only cells with series resistance smaller than 40MΩ were included in the dataset. Cells that required more than 200 pA of holding current to maintain these potentials were excluded from the data set.

### Western blot

Protein samples were denatured by boiling in SDS sample buffer containing 10% β-mercaptoethanol and then subjected to 10% SDS-PAGE. Blots were transferred onto a nitrocellulose membrane and blocked with Blotting Grade Blocker (#170-6404, Bio-Rad, Hercules, CA). Proteins were transferred to a nitrocellulose membrane, then immunoblotted with appropriate primary antibodies. Membranes were incubated overnight with the following primary antibodies: p-eIF2α (1:1000; #3398, Cell Signaling, Danvers, MA); total eIF2α (1:1000; #2103, Cell Signaling); Tubulin (1:40,000; #SAB4500087, Sigma-Aldrich) Puromycin (1:10,000; #MABE342, Millipore, Burlington, MA); and ß-actin (1:12,000; #sc1616, Santa Cruz Biotechnology, Dallas, TX) in 5% BSA in TBST at 4°C and subsequently washed thrice for 5 minutes with TBST. Incubation with the HRP-conjugated secondary antibodies goat anti-rabbit, goat anti-mouse or rabbit anti-goat (1:10,000; Jackson Immunoresearch, West Grove, PA) was done for 1 hour at room temperature. Immunodetection was carried out with EZ-ECL kit (#20500120, Biological Industries, Beit Haemek, Israel), acquired with Chemi XRS Gel Doc (Bio-Rad), and quantified using the Quantity One software (Bio-Rad). Intensity was normalized to background signal.

### SUnSET

Cells were incubated with 1µg/mL puromycin for 10 minutes and harvested for either western blot or fixed for immunodetection. For western blot, cells were lysed using puromycin lysis buffer (PLB; Tris 40 mM, NaCL 0.15 M, ß-glycerol 25 mM, NaF 50 mM, Na_3_VO_4_ 1 mM, glycerol 10%, Triton x100 1%) and diluted 1:1 with SDS sample buffer containing 10% β-mercaptoethanol.

### Immunocytochemistry

Cells were washed with PBS and fixed in 4% paraformaldehyde in PBS for 10 minutes. Cells were then permeabilized with 1% Triton X-100 (in PBS; PBS-T 1%) thrice for 5 minutes and blocked for one hour in blocking buffer (10% FBS, 0.3% BSA, and 1% Triton X-100 in PBS) at room temperature. Primary antibodies in blocking buffer were then added, and incubated overnight at 4°C. Next, cells were washed thrice for 5 minutes in PBS-T 1%, and incubated with secondary antibodies 1:500 for 1 hour at room temperature. The following primary antibodies were used: αchicken Map2 (1:1000; #ab5392, Abcam, Cambridge, UK) or αmouse Map2 (1:250; #M9942, Sigma-Aldrich), GFAP (1:2000; #G3893, Sigma-Aldrich), Calreticulin (1:500; #315003, Synaptic Systems, Goettingen, Germany). Secondary antibodies used: Alexa Fluor goat anti-chicken 488 (#ab150169), 568 (#ab175477), and 647(#ab150175) and goat anti-mouse 647 (#ab150119-500) (all from Abcam), as well as goat anti-mouse 488 (#A-11001, ThermoFisher Scientific). Prior to mounting, cells were washed thrice for 5 minutes with PBS-T 1% and twice with PBS for 5 minutes. Cells were mounted with ProLong Diamond Anti-Fade Mountant with DAPI (#P36962, ThermoFisher Scientific) on pre-cleaned 26×76 mm microscope slides of 1.0-1.2 mm thickness (#BN1045001CG, Bar Naor, Petah Tikvah, Israel).

### Fluorescent microscopy

#### Image acquisition

Cortical primary cultures were imaged at x60 magnification with Olympus IX 81 microscope using cellSens Dimensions software (Olympus, Shinjuku, Japan). 1 pixel=108 nm. Images were acquired in z-stacks of 0.25 µm steps.

#### tRNA distribution analysis

Distribution of labeled tRNA was quantified using Fiji software and images of (1) Map2/GFAP (and Calreticulin for co-localization analyses) positive cells and (2) Cy3-labeled tRNA by converting images to binary mask (black and white) and superimposing them for points of overlap.

#### FRET acquisition and analysis

FRET signal was captured using the following complete filter sets Cy3 (donor; Chroma 49004) and Cy5 (acceptor; Chroma 49006). FRET analysis was performed by measuring the mean intensity of particle signals in each of these channels: Cy3, Cy5, and Youvan (cellSens FRET module). FRET signals were calculated after subtracting the bleed-through and cross-excitation fluorescence measured in control wells containing cells transfected with either Cy3- or Cy5-labeled tRNAs and are presented as calculated FRET (cFRET). Control wells were also stained with Map2-Alexa 488 to account for possible bleed-through to cy3 channel. Intensity and particle area were determined using FIJI Image J software. Total intensity was calculated as particle area X intensity. FRET intensity was calculated as follows: 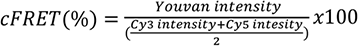.

tRNA localization was calculated as a function of distance from the nucleus using the MATLAB-based algorithm SynD (96), which provides automated analysis of immunofluorescence images of antibody-labeled neurons. The program produces an overlay of neuron morphology and detects differently labeled desired molecules within it.

#### Live imaging

Live primary cultures were imaged in an environmental chamber at a constant temperature of 37°C containing 5% CO_2_ to maintain cell viability in MEM medium without phenol red (#51200046, ThermoFisher Scientific), to allow for optimal visibility. Thirty frames were collected at 1 min intervals. Particle tracking and motion analysis were performed using the cellSens Dimension module. Prior to imaging, axons were stained with Alexa Fluor 488-conjugated cholera toxin (#C22841, ThermoFisher Scientific). For kymograph analysis, images were acquired at 6 sec intervals, and transport was determined in FIJI Image J software using the Kymo ToolBox plugin (102).

### Super resolution microscopy

#### Image acquisition

The localization accuracy of the optical system was determined from the localization of centroids of point spread functions generated by 100 nm fluorescent beads, which were imaged for 4000 frames at similar laser intensities recorded for the dSTORM measurements on immunolabeled samples (Figure 4 – figure supplement 1 A,B). The centroid of the lateral spread of the fluorescence intensity was calculated for every frame and an accuracy of detection histogram was obtained for all the localizations in time (Figure 4 – figure supplement 1A). The resulting spread was modeled using a bi-dimensional Gaussian function providing localization precision (σ) of ∼19.4 nm (Figure 4 – figure supplement 1C). The full width at half maximum was calculated to be ∼44.6 nm (Figure 4 – figure supplement 1D). Graphs and the related statistics were produced using MATLAB Statistics Toolbox Version 10 of MATLAB R2015a (The MathWorks, Natick, MA).

#### DNA–antibody conjugation

Amine groups of the lysine residues from secondary antibodies (#715-005-151 and #711-005-152, Jackson ImmunoResearch; for more detail see Table S1) were modified with maleimide–PEG_2_–NHS ester (Sigma Aldrich) in PBS (pH 7.4) for 2 hours at room temperature and purified using Zeba spin desalting column (7000 Da MWCO; #89883, ThermoFisher Scientific). Thiolated DNA oligos (Integrated DNA Technologies, Coralville, IA; see Table S2) were reduced using DTT. Excess DTT was removed using NAP-5 column (GE Healthcare, Chicago, IL) with PBS (pH 7.4) as eluent. Maleimide conjugated antibodies and thiol terminated oligonucleotides were mixed in a microcentrifuge tube and stirred at 4°C for 12 hours. Finally, DNA conjugated antibodies were purified and concentrated using Amicon Ultra Centrifugal Filter (100 kDa MWCO) and were used for secondary immunostaining in 1:300 dilution.

The complementary oligonucleotides were conjugated with Atto 655 NHS ester (Atto technology, Amherst, NY) in 0.1 M NaHCO_3_ and purified by reversed phase high performance liquid chromatography (HPLC). Fluorophore-conjugated oligos were characterized by matrix-assisted laser desorption ionization mass spectrometry (MALDI–MS) (Table S3) (104). For more detailed protocols see Supplementary Information.

#### Single molecule localization microscopy and image analysis

Samples were imaged at 37°C in a closed chamber (Ludin Chamber, OKO lab, Pozzouli, Italy) specialized for loading 18 mm round coverslips and mounted on an inverted motorized microscope (Olympus IX83, Olympus) equipped with a 100X1.49NA PL-APO objective and an azimuthal drift control device, allowing long acquisition in oblique illumination mode using multi-laser (namely, 405 nm, 491 nm, 532 nm, 561 nm, and 640 nm) launch (Roper Scientific, Vianen, Netherlands). The images were acquired at the center quadrant (256X256 Pixel^2^) of an EMCCD camera (Evolve, Photometrics, Tucson, AZ). The illumination and acquisition was controlled by MetaMorph version 7.8 (Molecular Devices). dSTORM Imaging was performed in an extracellular solution containing a reducing and oxygen scavenging system, according to the dSTORM (86). Prior to imaging, a second fixation was performed with 4% paraformaldehyde. Custom synthesized gold nanoparticles (100 nm diameter) were used as fiduciary markers. The immunolabeled cells were imaged in a dSTORM buffer with a cocktail of chemicals (see supplementary information) to induce stochastic activation of sparse subsets of molecules. dSTORM buffer was added prior to imaging. Photoconversion of carbocyanine dyes from ensemble to single molecule density was achieved by illuminating the sample with 300 mW (laser power measured at the source) excitation laser to convert the fluorescence molecules into the metastable dark state. The fluorescence emission above 640 nm was diverted to the camera port via a quad-band dichroic mirror (Part No. FF01-440/521/607/700-32, Semrock, Rochester, NY).

After achieving an optimal density of 0.01-0.04 molecules/µm^2^ the illumination laser power was optimized to 150 mW and 5 stacks of 4000 frames each were obtained with an exposure time of 20ms per stack, acquiring a total of 20,000 images. The instantaneous density of single molecules was controlled by modulating the power of 405 nm laser.

For DNA Exchange PAINT, for imaging rabbit rpS6, corresponding imager strand (2.5 nM) conjugated to ATTO 655 was diluted in PBS in 1:1000. For imaging mouse rpL10a, imager strand (100 nM) conjugated to ATTO 655 was diluted in PBS in 1:100. Imaging was done sequentially at 100 to 200 ms frame rate separated by three times of five minute washes with PBS to clear residual imager strand. The laser power used was 300 mW at the source. Subsequent image processing and analysis were performed using ThunderSTORM (93), an ImageJ plugin to reconstruct dSTORM images. Images were then exported to Metamoprh for representation of pseudocolor overlay analysis.

Overlap analysis was done by thresholding all images at the same fixed point in a far red channel (minimal background autofluorescence), using a threshold that captures signals between 2 and 255 in an 8 bit image. Using Image Calculator of ImageJ, fractional overlap was calculated according to the following formulae:

Formula for calculating two color overlap = (R*G)/(R + G – (R*G))

Formula for calculating three color overlap = (R*G*B)/[R + G + B – (R*B) – (R*G) – (G*B) + (R*G*B)]

R = thresholded rpS6 super resolution image, G = thresholded rpL10a super resolution image, B = thresholded tRNA super resolution image

SR Tesseler (Levet et al., 2015) was used to segment the output list of localizations generated by ThunderSTORM. For the rpL10a and rpS6 signal, two distinct peaks of densities were identified and the localizations were segregated based on the bimodal distribution. In the next step, overlap analysis as described above was performed on different distributions of rpL10a and rpS6 (Figure 4 – figure supplement 1).

For characterizing tRNA puncta, ROIs were created around the tRNA puncta and their intensities and areas were determined using FIJI. The ROIs were transferred to thresholded images of rpL10a and rpS6 to determine which of the ROIs contain non-zero intensities. These qualified as overlapping tRNA puncta, and their areas were selected for distribution analysis.

### Chemical LTP

Chemical LTP was induced in culture using 30 µM bicuculline (Bic, #0130, Tocris Bioscience, Bristol, UK) and 100 µM 4-Aminopyridine (4-AP, #275875, Sigma Aldrich). GluR inhibition was performed using 40 µM D(-)-2-amino-5-phosphonopentanoic acid (APV, #A8054, Sigma Aldrich) and 2 µM 6,7-dinitroquinoxaline-2,3-dione (DNQX, #D0540, Sigma Aldrich).

### Statistical analysis

Statistical analysis was performed using GraphPad Prism 8.0.1 for windows, GraphPad software, San Diego, CA (www.graphpad.com). Normality was determined using the Shapiro-Wilk test, F-test to compare variances, and/or Bartlett’s test to compare SDs. Outliers were detected using ROUT test (Q=1%). For two-group comparisons, Student’s t-test was used for Gaussian distribution and Mann-Whitney test for non-normal distribution. For multiple comparisons, one-way ANOVA followed by Dunnett’s multiple comparisons test with Tukey’s post hoc test was applied to Gaussian distributions with normal distribution of SDs, Brown-Forsythe and Welch’s ANOVA was used for Gaussian distributions with unequal SDs, and Kruskall-Wallis test followed by Dunn’s multiple comparisons test was applied to non-normal distributions with unequal SDs. For grouped analyses, either two-way ANOVA was applied or Holm-Sidak multiple t tests. Significance was set at p≤0.05.

## Results

### Neurons transfected with fluorescently-labeled tRNA are healthy and functional

We first examined whether introduction of fluorescently-labeled tRNA adversely affects neuronal health or function, as previous work with fluorescently-labeled tRNA has thus far been limited to somatic cells and cell lines (8, 55). For this purpose, fluorescently-labeled tRNA was introduced into mouse-derived primary cortical cultures using a standard transfection protocol (schematic summary in Figure 1A, see also Methods). After transfection was attempted with several different reagents, PolyJet transfection reagent was found to yield the highest transfection efficiency (approximately 80% transfected cells) with the minimum required reagent concentration (1 µl PolyJet and 100 ng tRNA for 87,000 cells). Cell viability and morphology were maintained post-application of labeled tRNA (Figure 1B). These findings were further supported by measurements of expression levels of phosphorylated α-subunit of eukaryotic initiation factor 2 (eIF2α). Phosphorylation of eIF2α is a direct result of environmental changes and/or stress, and leads to suppression of general protein synthesis (85, 101). Levels of phosphorylated eIF2α returned to their basal state within 48 hours following transfection (Figure 1 – figure supplement 1). Based on these findings, all experiments were conducted a minimum of 48 hours following transfection.

**Figure 1.**
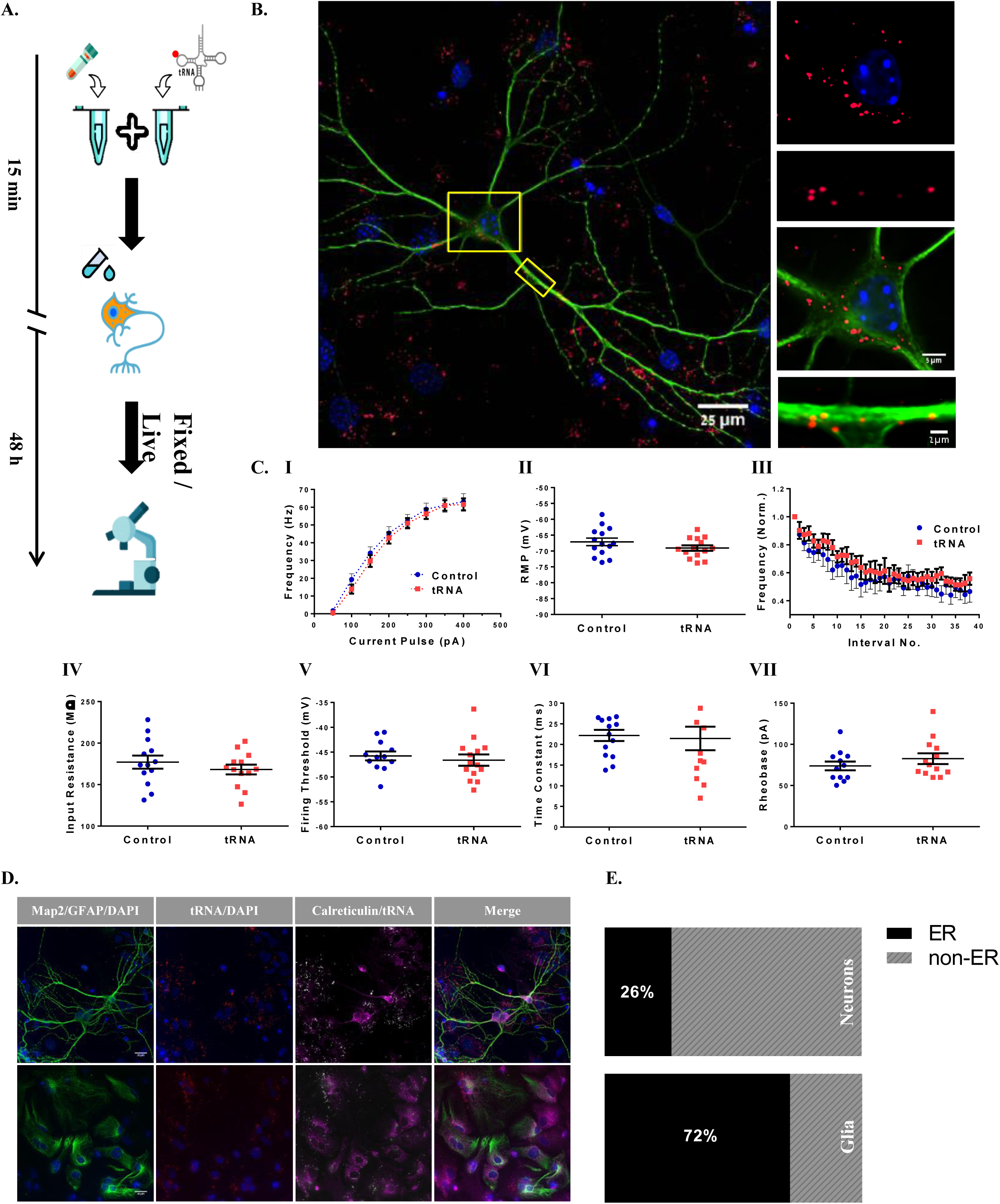
Differential distribution of tRNA in neurons and glial cells in cortical primary culture. **A-C. Neurons in cortical co-culture transfected with labeled tRNA maintain function and morphology. A. Schematic illustration of introducing fluorescently labeled tRNA to cortical primary culture**. Labeled tRNA (4nM) was mixed with a transfection reagent (PolyJet^TM^) and dripped onto cells after a short incubation period (15 min). tRNA distribution was viewed after 48 hours under a microscope. **B. Viable neuronal culture transfected with labeled tRNA.** Representative images of a neuronal-glial co-culture stained with Map2 antibody with magnification of (I) neuronal processes and (II) somata, demonstrating fluorescently-labeled tRNA. **C. Basic neuronal intrinsic properties are maintained following application of labeled tRNA.** Several active and passive electrophysiological intrinsic properties were measured in brain slices transfected with fluorescently-labeled tRNA. No differences were observed between tRNA treated and control neurons in any of the properties examined. (I) Firing frequency during 500ms-long pulses of depolarizing current injections; (II) RMP – resting membrane potential; (III) Normalized instantaneous firing frequencies as a function of interspike interval for the first 38 interspike intervals; (IV) Input resistance; (V) Firing threshold; (VI) Membrane time constant; (VII) Rheobase. n(tRNA treated cells)=14, n(control cells)=14 (both obtained from 3 mice). Data are represented as Mean ± SEM (p>0.05). **The majority of labeled tRNA is co-localized with rER in glial cells but not in neurons in mouse primary co-culture. D. Representative immunocytochemical staining for neurons and glial cells.** Immunostaining of neurons (Map2; I-IV) and glial cells (GFAP; V-VIII) with Calreticulin, a marker for rough ER **E.** Quantification of data collected from images in (D), presented as bar diagrams.

To further rule out any effects of the labeled tRNA on neuronal function, we tested the effect of the maximal amount of tRNA that could enter a single cell during transfection (100 ng) on the intrinsic neuronal properties of the CA1 region of the hippocampus in mouse brain slices using the patch-clamp technique (Figure 1C). Properties tested included firing frequency, recovery period, resting membrane potential, adaptation ratio, input resistance, firing threshold, time constant, and rheobase. None of the properties tested showed any difference between the tRNA-patched cells and control cells (patched only with K-gluconate internal solution), suggesting that the introduction of the labeled tRNA at our working concentration does not cause neuronal stress that can be measured electro-physiologically.

As tRNA is a key player in protein synthesis, we next tested its functionality by analyzing the co-localization between fluorescently-labeled tRNA and the central site of protein synthesis in neurons as well as in glial cells: the rough endoplasmic reticulum (rER) (Figure 1D). As expected, the majority (72%) of the labeled tRNA in glia was found to be co-localized with the rER protein Calreticulin, while in neurons only 26% of the tRNA co-localized with the rER (Figure 1E). Notably, although rER is present in proximal dendrites (47), we found labeled tRNA in distal dendrites unbound by rER, consistent with the notion of free ribosomes (4).

### Differential dynamics of tRNA motion and puncta size between neuronal sub-compartments and glial cells

After observing tRNA in fixed cultures, we next characterized tRNA in live cultures, focusing on motion (via mean instantaneous velocity of the tRNA puncta) and puncta size (measured as number of separate objects in a single track) parameters. Imaged cultures, containing neuronal networks on a surrounding layer of glial cells, were transfected with tRNA labeled with fluorescent dye Cy3, which has strong fluorescence and was stable throughout the imaging time frame (manifested as a consistent number of detected objects (Figure 2 – figure supplement 1)).

Thirty-minute films were acquired at one-minute intervals at four consecutive time points of representative culture fields. The baseline was assayed at t0. At t1 puromycin was applied and at t2 washed out. t3 was acquired 90 minutes after puromycin application in order to test a return to basal levels. Mean instantaneous velocity of the particles was derived from the displacement of puncta between each acquisition. Our results suggest that mean instantaneous velocities of tRNA distribute non-normally in the examined locations, with the median mean instantaneous velocities in the neuronal soma and glial cells being relatively similar at 22.16 nm/s (inter quartile range (IQR) =18.28-26.02 nm/s, Figure 2A I) and 20.77 nm/s (IQR=17.35-23.68 nm/s, Figure 2A III), respectively. In dendrites, the median mean instantaneous velocity observed was significantly lower at 15.49 nm/s (IQR=11.1-19.86 nm/s, p<0.0001, Figure 2A II). Importantly, our observations fall in line with previous data regarding RNA, and specifically mRNA kinetics in neurons, which suggests that the average velocity of total mRNA in neurons is approximately 1-100 nm/s (49, 50). Addition of the translation inhibitor puromycin [50µM] had no significant effect on tRNA velocity in the soma or dendrites, and no change was observed over time.

**Figure 2.**
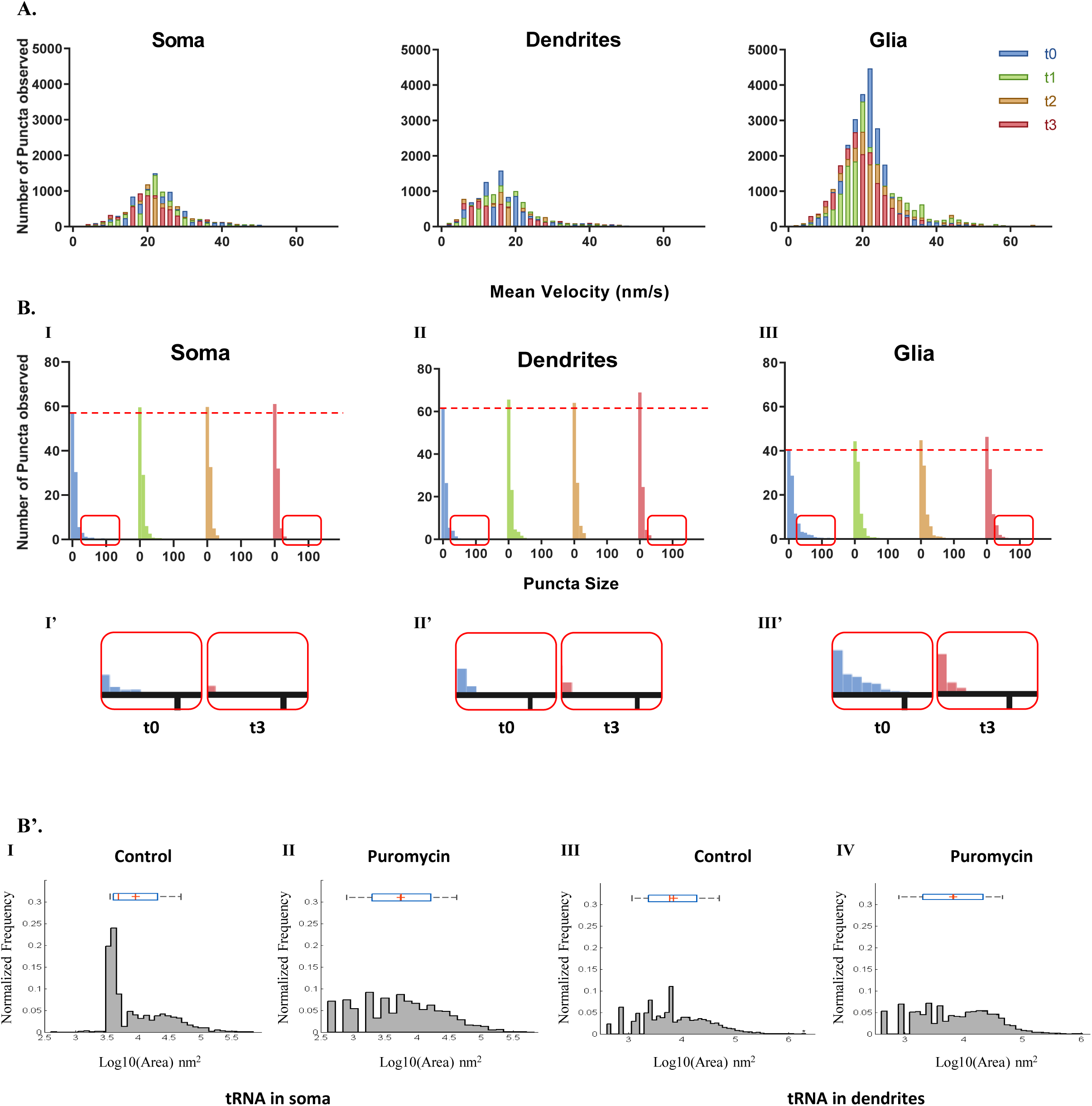
tRNA velocity in live primary cortical culture indicates slow transport rather than free diffusion as well as differential dynamics in somata and neuronal processes. **A. tRNA velocity in dendrites and axons is lower compared to somata and glia.** Histograms of mean velocities (nm/s) of tRNA puncta in (I) somata, (II) dendrites, and (III) glial cells, measured at 4 time points within constant ROIs in 3 representative monitored frames. t0=0-30 min, t1=30-60 min, t2=60-90 min, t3=90-120 min. Puromycin added at t1 and washed out at t2. **B. tRNA puncta dissociate following puromycin treatment as measured by light microscopy.** Histograms of puncta size frequencies (measured as multiplications of the smallest detectable signal, termed signal units) in (I) somata, (II) dendrites, and (III) glial cells, measured at 4 time points within constant ROIs in 3 representative monitored frames. t0=0-30 min, t1=30-60 min, t2=60-90 min, t3=90-120 min. Puromycin added at t1 and washed out at t2. Dotted line marks the number of puncta ≤2 signal units at t0 at each examined compartment (I’-III’). Magnifications of puncta size frequencies ≥8 signal units at t0 and t3 in somata, dendrites and glial cells, respectively. **B’. tRNA puncta dissociate following puromycin treatment as measured by direct STochastic Optical Reconstruction Microscopy (dSTORM).** (I-II) Histograms of tRNA puncta in somata (I) under basal conditions; Mean=9120.1nm^2^, SD=2.95nm^2^, median=4786.3nm^2^, IQR 3981.1nm^2^-19952.6nm^2^; (II) following puromycin addition; Mean=5623.4nm^2^, SD=4.4nm^2^, median=5623.4nm^2^, IQR 1995.3nm^2^-15848.9nm^2^; (III-IV) Histograms of tRNA puncta in dendrites (III) under basal conditions; Mean=7079.5nm^2^, SD=4.07nm^2^, median=6025.6nm^2^, IQR 2398.8nm^2^-19054.6nm^2^; following puromycin addition; Mean=6606.9nm^2^, SD=4.6nm^2^, median=6760.8nm^2^, IQR 1995nm^2^-19952.6nm^2^.

tRNA puncta sizes were highly variable. At t0, the majority of labeled tRNA was found in small puncta (≤6 objects), however, many observed puncta measured over 100 objects in size, especially around the nuclear surface. These puncta may represent polyribosomes anchored to the rER. In order to interfere with the basal level of mRNA translation in the culture, puromycin was added to the culture for 30 minutes and then washed out. Puromycin causes ribosome dissociation during elongation (48), and therefore we hypothesized that puromycin would dissociate the assembled ribosome from the rER, thus increasing the free ribosome fraction (48). In line with our hypothesis, puncta size distribution in all locations was drastically altered immediately following addition of puromycin, as the number of larger puncta decreased, while the number of smaller puncta increased (soma: p=0.0243; dendrites and glia: p<0.0001; Figure 2B I-III, with magnification of the disappearance of the “tail” of larger puncta in I’-III’). This effect persisted after puromycin washout, possibly due to disassembly of the ribosomal complex. Importantly, the total number of the objects remained unchanged throughout the experiment (Figure 2 – figure supplement 1), indicating that the total number of labeled tRNA molecules remained constant while their distribution changed.

To further verify these findings, we examined the dynamics of tRNA puncta using super-resolution microscopy, separately monitoring tRNA puncta in somata and dendrites. This revealed that addition of puromycin led to a decrease in puncta size (determined by area), due to dissociation into smaller puncta, as evident by the increase in the number of smaller puncta observed (Figure 2B’). These findings are similar to the results obtained using fluorescent microscopy, with a more pronounced effect visible in somata than in dendrites. Super-resolution microscopy also corroborated our findings that the basal distribution of puncta size in dendrites leaned towards smaller puncta compared to the soma and to glia cells.

### tRNA motility in dendrites

In order to study tRNA motion within dendrites with finer temporal resolution, Cy3-labeled tRNA was monitored in a neuronal co-culture stained with neuron-specific dye NeuO (see Methods) over a period of 10 minutes, acquiring images in 6-second intervals (Figure 3A). A kymograph showing movement along a single axis of a representative film shows the path of movement of three distinct tRNA puncta (Figure 3B). We performed directional analysis in order to characterize the dendritic tRNA movement. Two of the dendritic tRNA puncta appeared to alternate between the anterograde and retrograde directions, thus moving a larger overall distance than the final displacement between the start and end points, while a third puncta remained static throughout the film (Figure 3C).

**Figure 3.**
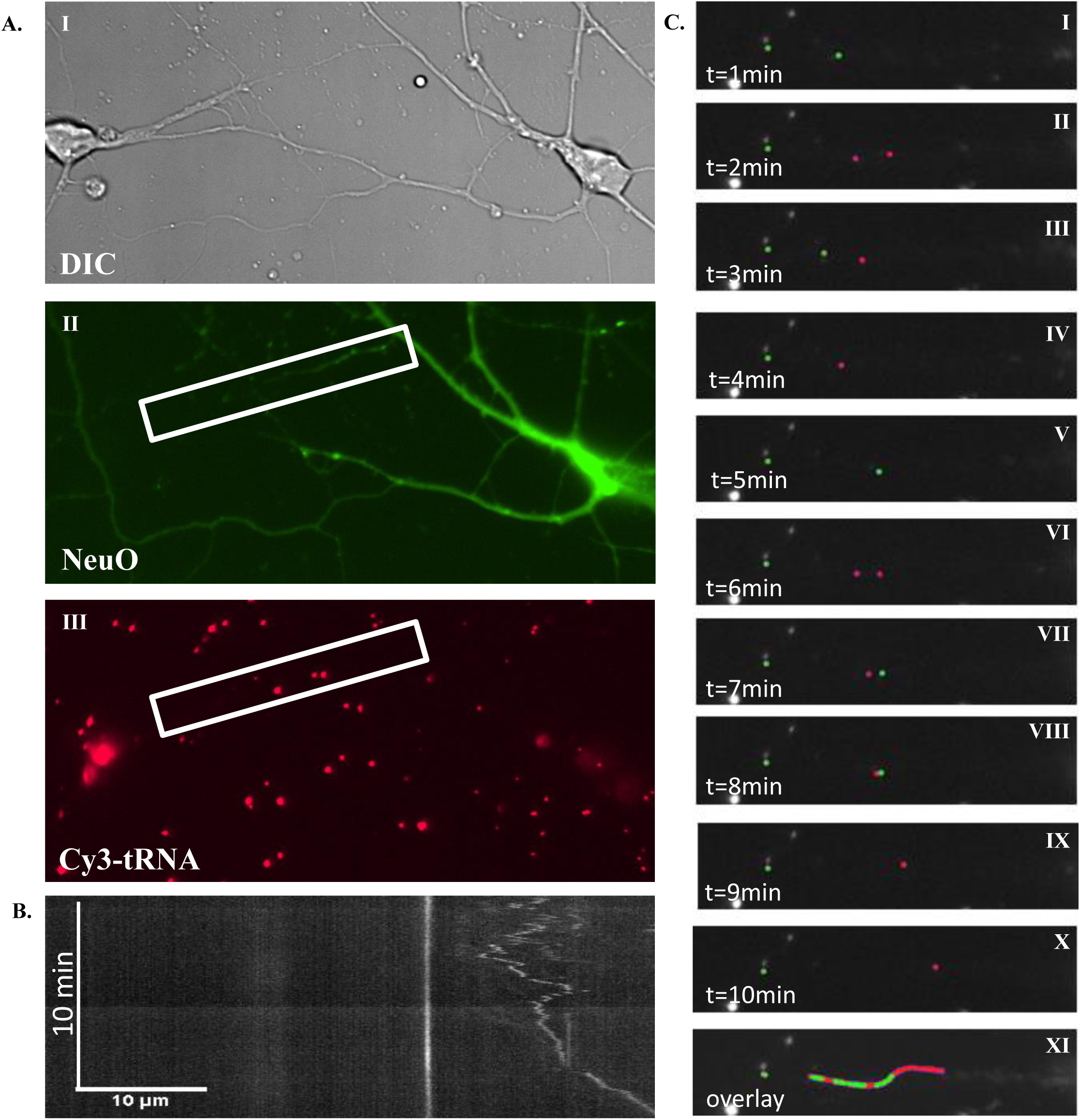
tRNA is bi-directionally transported in dendrites. **A. Still image capture of live neurons.** Images of neuronal culture stained with NeuO NeuroFluor live neuronal dye and transfected with tRNA labeled with Cy3 (I-III). **B. tRNA is transported in dendrites.** Kymograph of tRNA movement within dendrites over a 10-minute period. Images acquired at 6-second intervals. **C. tRNA in dendrites is transported in both anterograde and retrograde directions.** Representative captures of tRNA movement in 1-minute intervals over a period of 10 minutes. The different directions are marked in different colors. tRNA molecules are represented in red (I-X). (XI) Overlay of images I-X.

### The assembled ribosome co-localizes with fluorescently-labeled tRNA

Protein synthesis occurs when aminoacyl-tRNA is associated with both ribosomal subunits. To establish the functionality of the labeled tRNA, we examined its co-localization with each of the ribosomal subunits and with the assembled ribosome in cultured neurons. Using dSTORM, we were able to visualize nanoscale organization of the small ribosomal subunit protein rpS6, the large ribosomal subunit protein rpL10a, and of tRNA (Figure 4A). We differentiated between clustered and diffused fractions of each of the proteins, rpS6 and rpL10a, based on the density of localization, defining the threshold as 40 localizations/nm^2^. Both diffused populations responded similarly to puromycin addition, showing a reduction in the size of the diffuse fraction (Fig 4B). We next measured the co-localization of the clustered and diffuse fractions with the two ribosomal proteins (Figure 4C I-VI). To test the functionality of each fraction, we also measured co-localization of rpS6 and rpL10a after puromycin addition. We found that despite a high level of co-localization between the clustered pool of rpS6 and diffuse pool of rpL10a, and vice versa, this distribution was unaffected by puromycin addition (Figure 4C VII), pointing to the possibility of random overlap between components considered as part of the diffuse pool. The co-localization of the clustered pool of rpS6 with the clustered pool of rpL10a was minimal (<5%) and also unresponsive to puromycin, whereas approximately 45% of the diffuse pools of rpS6 and rpL10a co-localized (Figure 4C VIII). Notably, the co-localization of the diffuse pools decreased significantly following addition of puromycin to approximately 27%±3% (p=0.0053, Figure 4C VIII). These data suggest that the active ribosome complex consists of the diffuse fractions of both ribosomal proteins and disassembles upon puromycin application, consistent with the decreased overlap observed.

**Figure 4.**
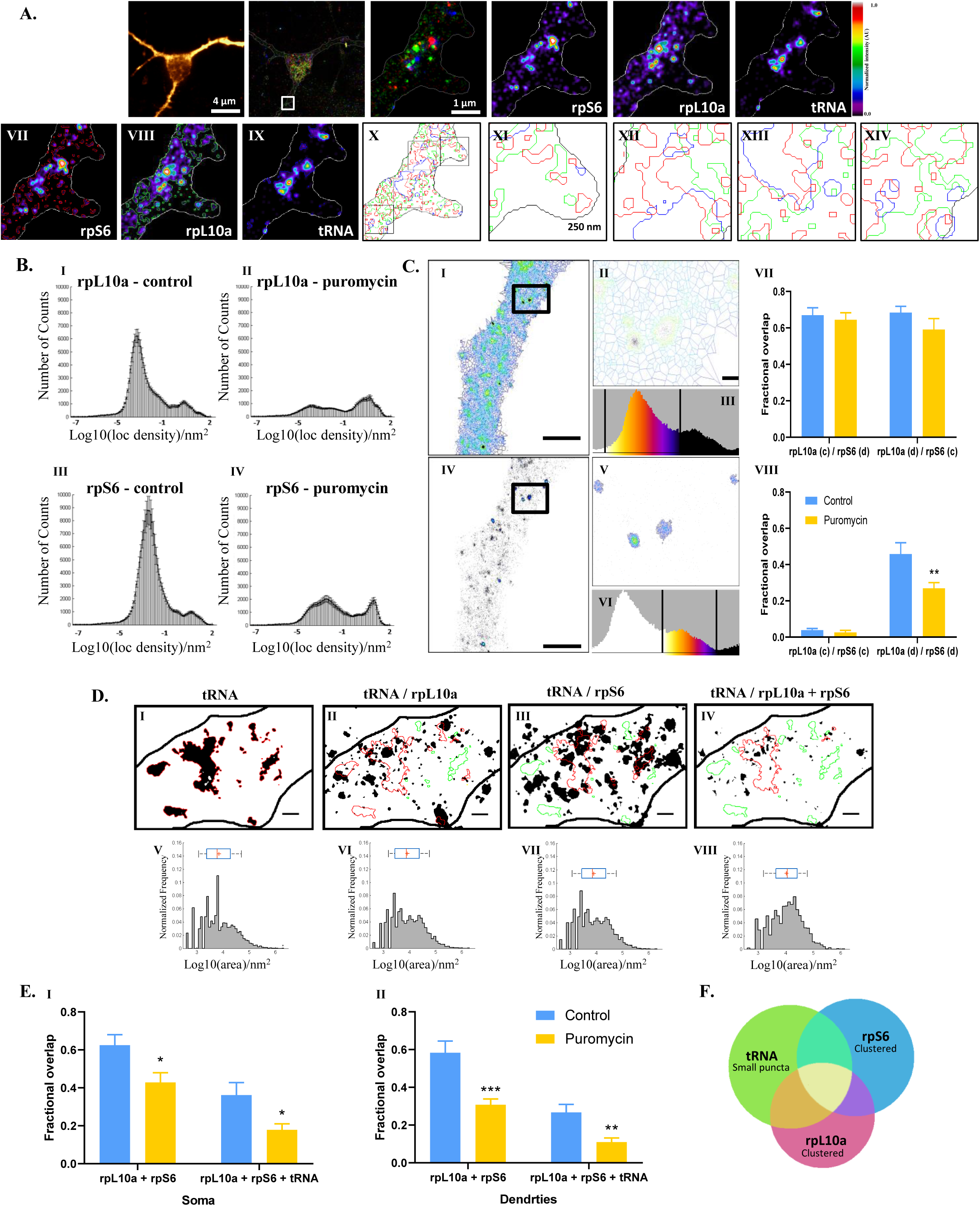
A fraction of fluorescently labeled tRNA is associated with the assembled ribosome as measured in neurons by single molecule based localization microscopy. **A. Nanoscale organization of translation components - rpL10a, rpS6, and tRNA.** (I) Representative image of a Map2-positive neuron; (II) 3 color single molecule localization microscopy of rpS6 (red), rpL10a (green), and tRNA (blue); (III) magnification of region of interest in a dendrite; (IV) rpS6 in ROI; rpL10a in ROI; (VI) tRNA in ROI; (VII-IX) Semi-automated detection of signals differently outlined for each staining of rpS6, rpL10a, and tRNA, respectively; (X) 3 color overlay of the segments obtained in VII-IX; (XI-XIV) magnification of 4 regions of interest in picture X. **B. Puromycin treatment leads to a decrease in diffuse pools of rpL10a and rpS6 based on localization density.** (I) Total number of counts of rpL10a in control = 1268723. (II) Total number of counts of rpL10a in puromycin = 536530. (III) Total number of counts of rpS6 in control = 2056885. (IV) Total number of counts of rpS6 in puromycin = 765686. Loc=localizations **C. Puromycin treatment leads to decreased overlap of diffuse pools of L10a and rpS6 but not in others.** Diffuse (I-II) and clustered (IV-V) regions of rpL10a labeling in a representative dendrite segmented based on the distribution of density of localizations (III and VI, respectively). Vertical lines define the boundaries of each region. Scale bar: 2 µm, 250 nm. (VII-VIII) Fractional overlap between (VII) clustered rpL10a and diffuse rpS6, clustered rpS6 and diffuse rpL10a, and (VIII) clustered rpS6 and rpL10a, and diffuse rpS6 and rpL10a under basal conditions and after application of puromycin. n=15 different captured regions from 3 biologically independent repeats. ** p<0.01 **D. Larger tRNA puncta more commonly associate with L10a and rpS6 than smaller puncta.** (I) ROIs of thresholded tRNA puncta. (II-IV) Transfer of ROIs from I to rpL10a, rpS6, and the overlapping regions of rpL10a and rpS6, respectively. Red=overlap, green=no overlap. Scale bar: 500nm. (V) Logarithmic scale plot of total area of tRNA. Mean=7079.5nm^2^, SD=4.07nm^2^, median=6025.6nm^2^, IQR 2398.8nm^2^-19054.6nm^2^; (VI-IX) Logarithmic scale plots of total area of overlap in II-IV, respectively. (VI) Mean=8317.6nm^2^, SD=4.07nm^2^, median=7585.8nm^2^, IQR 2818.4nm^2^-25118.8nm^2^; (VII) Mean=7943.3nm^2^, SD=4.17nm^2^, median=7244.4nm^2^, IQR 2754.2nm^2^-22387.2nm^2^; (VIII) Mean=10715.2nm^2^, SD=3.72nm^2^, median=11481.5nm^2^, IQR 3981.1nm^2^-25118.9nm^2^. **E. Puromycin treatment reduces the association of rpL10a, rpS6, and tRNA.** Fractional overlap of rpL10a and rpS6 alone and together with tRNA puncta in (I) somata and (II) dendrites under basal conditions and following puromycin application. n=22 different captured regions (somata) and 14 different captured regions (dendrites) from 3 biologically independent repeats. * p<0.05; ** p<0.01; *** p<0.001 **F. A schematic summary of results.** Diffuse pools of rpL10a and rpS6 are found to overlap with each other as well as with larger tRNA puncta, whereas clustered regions of rpL10a and rpS6 are not found to co-localize, as well as small tRNA puncta.

Following characterization of the different fractions of representative ribosomal proteins, we proceeded to measure the degree of overlap between each element with labeled tRNA, separately and together (Figure 4D). Of the total tRNA population, 73% co-localized with rpL10a, whose median area was 7585.8nm^2^; 75% co-localized with rpS6, whose median area was 7244.4nm^2^; and 40% co-localized with rpL10a and rpS6, whose median area was 11481.5nm^2^. Thus, the larger puncta of the labeled tRNA co-localized to a higher degree with each of the ribosomal proteins both individually and mutually.

Addition of puromycin led to a significant reduction (approximately 40%) in the level of overlap between rpL10a and rpS6 (p=0.0168) and among rpL10a, rpS6, and tRNA (p=0.0193) in neuronal somata (Figure 4E I). An even more significant reduction (approximately 50%) was measured in the level of overlap between rpL10a and rpS6 (p=0.0008) and among rpL10a, rpS6, and tRNA (p=0.0045) in dendrites (Figure 4E II). Our findings regarding overlap between rpS6, rpL10a, and tRNA are schematically summarized in Figure 4F.

### mRNA translation as measured by FRET intensity is higher in young cultures

We next utilized labeled tRNA to measure FRET between Cy3- and Cy5-labeled tRNA, which allowed us to monitor active mRNA translation in primary murine cortical co-cultures (the principles of this method are visually demonstrated in Figure 5A and detailed in the Materials and Methods section of this paper). Images were acquired in the Cy3 (red), Cy5 (magenta), and raw FRET (yellow) channels, as well as DAPI and 488-Map2. After eliminating bleed through factors (calculated FRET; cFRET; white), we identified FRET puncta in fixed neurons.

**Figure 5.**
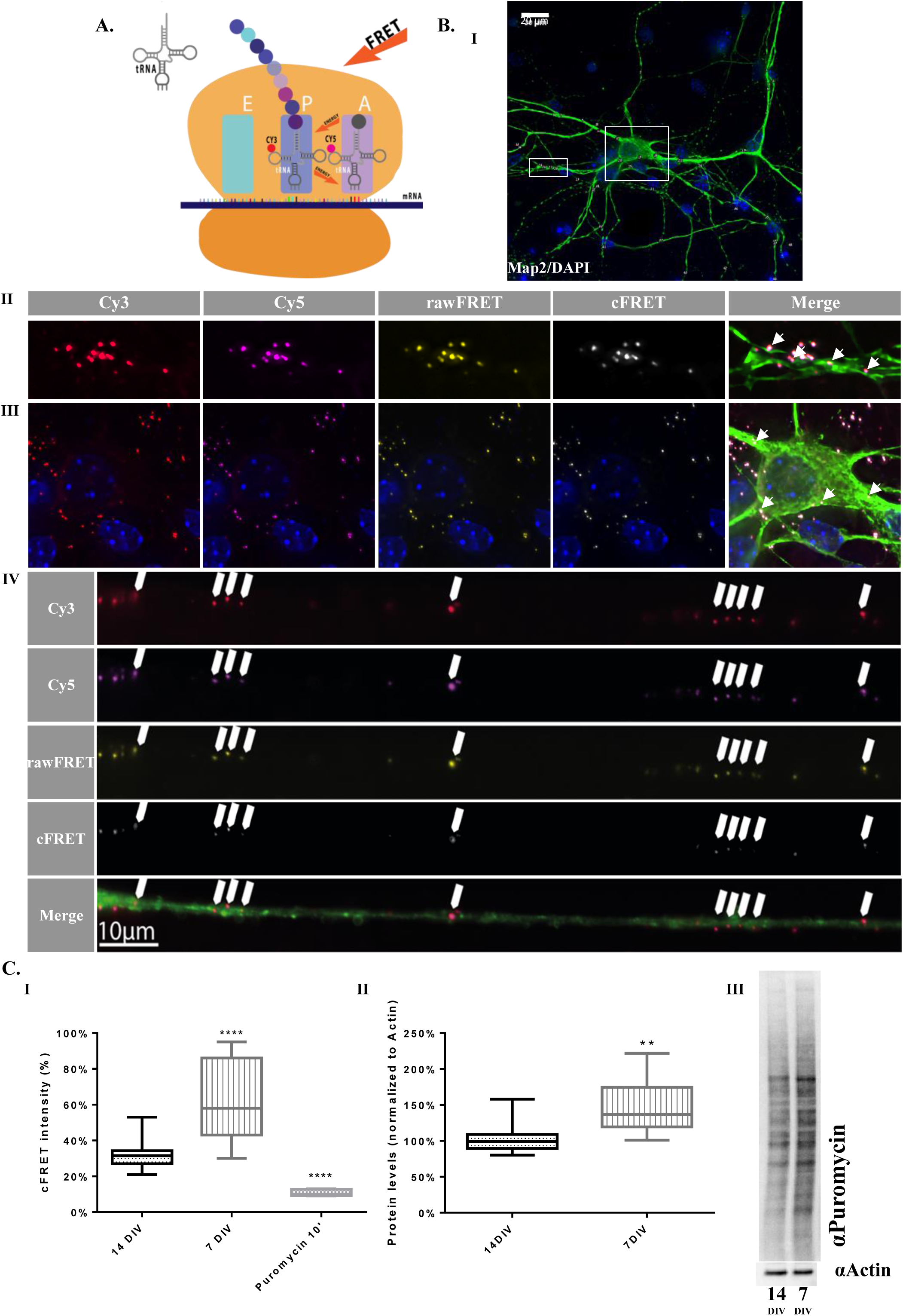
Levels of mRNA translation are negatively correlated with *ex vivo* developmental stage of neuronal primary culture. **A. Schematic diagram of monitoring active mRNA translation.** The FRET method utilizes tRNA pairs labeled as FRET donor (Cy3) and acceptor (Cy5) pairs and exploits the proximity of the tRNA molecules in the ribosomal A and P sites, as well as the temporal overlap, which are obligatory for the reaction to take place and enable monitoring active translation. **B. mRNA translation takes place throughout the neuron.** Representative image of (I) a Map2-positive neuron transfected with Cy3- and Cy5-labeled tRNA. (II) Magnification of a representative dendrite. (III) Magnification of the soma. (IV) Representative image of an axon infected with 488-CTX. White arrows indicate locations of active translation reported as FRET events. **C. mRNA translation levels are higher in young culture.** (I) cFRET signal intensity is higher in neurons in young culture (7 DIV) compared to mature culture (14 DIV). The signal was reduced below baseline after application of 50µM puromycin for 10 min. **** p<0.0001 (II) Puromycin incorporation levels are higher in young culture compared to mature culture. (III) Representative western blot of whole cell lysates of young and mature culture. ** p<0.01; n=3.

We next employed the tRNA-FRET method specifically in axons. To achieve this goal, cortical mouse neurons were seeded in microfluidic chambers as previously described, enabling the isolation of single axons (Figure 5 – figure supplement 1). The cells were transfected with tRNA, and 48 hours later live imaging was performed after incubation with fluorescently-tagged Cholera toxin (CTX) in order to visualize the axons (Figure 5B IV).

Younger, developing neurons exhibit higher levels of protein synthesis than mature neurons (51). Thus, we compared the global mRNA translation levels in neurons in young cultures and mature cultures (7 and 14 DIV, respectively) to further confirm the validity of our assay. FRET percentage in fixed neurons was calculated as the intensity of the signal normalized to area (Figure 5C). Our data suggest that global mRNA translation as measured by cFRET intensity was higher in young neurons than in mature ones (64%±4.7% and 32%±1.9%, respectively, p<0.0001; Figure 5C I). This was further validated by the reduction in global mRNA translation levels in mature neurons in response to puromycin, (11%±0.5% normalized cFRET intensity, p<0.0001). We further corroborated our results using the SUrface SEnsing of Translation (SUnSET) method, which showed approximately 1.5-fold up-regulation of global mRNA translation in young cultures, quite similar to the results obtained using the tRNA-FRET method (p=0.0081, Figure 5C II-III). As the SUnSET method requires lysed cultures, this result reflects the effect of glial cells present in the lysates, whereas with the tRNA-FRET method, we were able to measure mRNA translation specifically in neurons and within neuronal sub-compartments.

### cLTP induces biphasic mRNA translation up-regulation in neuronal somata and processes

We next employed the tRNA-FRET method to test the hypothesis that protein synthesis in both dendrites and neuronal somata is increased following chemical induction of long term potentiation (LTP), the main phenomenon of neuronal plasticity thought to underlie learning and memory (77–79, 105). LTP was chemically induced using potassium channel blocker 4-AP and GABA_A_ receptor antagonist Bicuculline (64). The experiments were carried out using primary cortical cultures derived from *Arc:dVenus* transgenic mice newborns (see Methods), providing a conclusive positive control of successful activation (Figure 6A). Levels of mRNA translation were measured at consecutive time points (baseline, 2, 4, and 6 hours post-activation), with spatial distinction between neuronal somata and dendrites. Somatic and dendritic mRNA translation were similarly affected at all-time points (ns, p>0.05), and revealed a biphasic pattern of up-regulation specifically 2 and 6 hours post activation (p<0.001 and p<0.0001, respectively; Figure 6B) with each sub-compartment individually producing the same biphasic pattern of up-regulation of mRNA translation (Figure 6B and Figure 6 – figure supplement 1). The similarity between somatic and dendritic mRNA translation levels was further demonstrated when presenting the ratio of cFRET intensity between somata and dendrites, which showed no difference at any of the time points examined (p=0.3115, Figure 6C).

**Figure 6.**
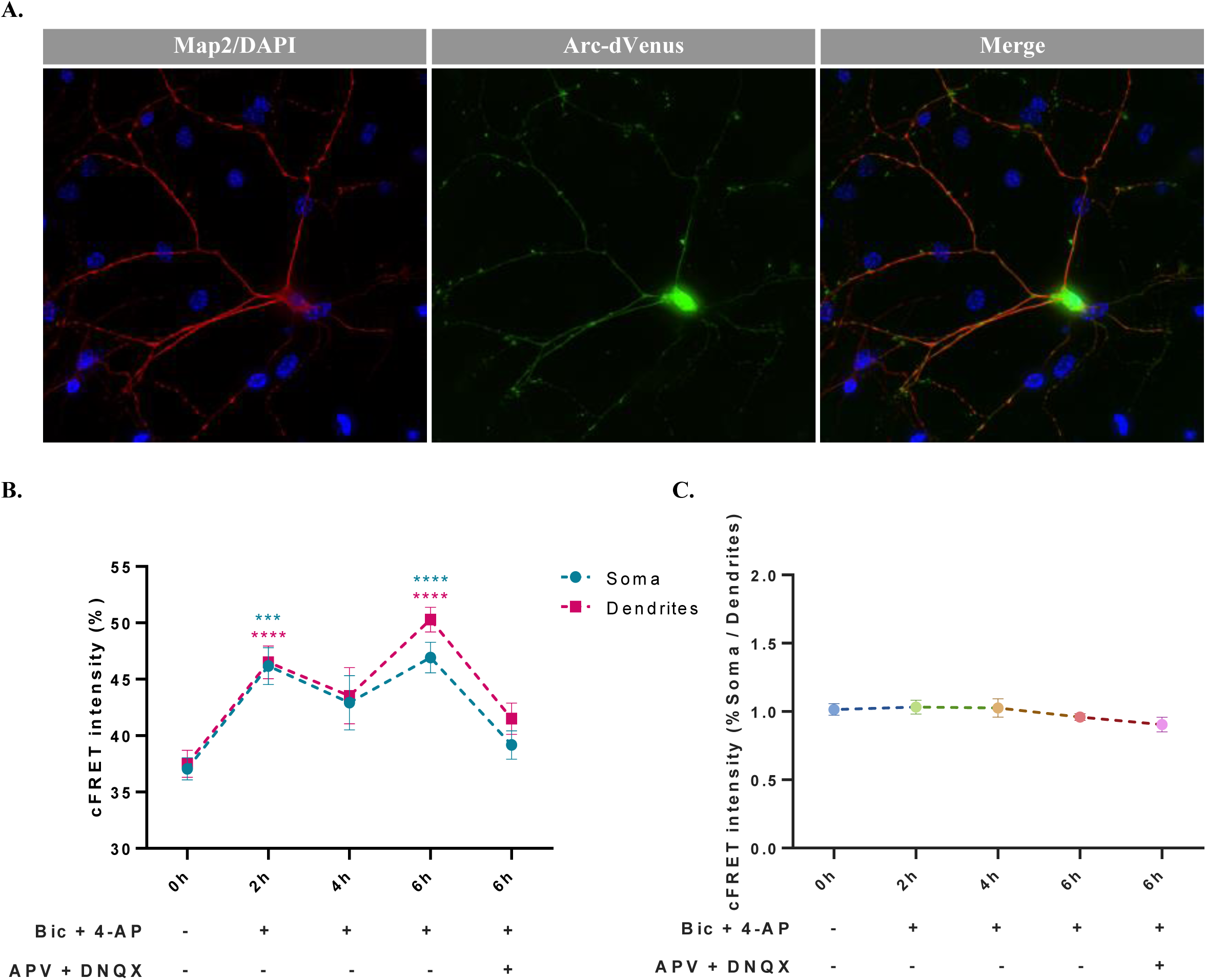
Biphasic mRNA translation up-regulation in neuronal processes and somata following cLTP. **A. Arc-dVenus detection after 6 hours reports successful induction of cLTP.** Representative images of an Arc-dVenus positive neuron stained with Map2 6 hours post activation. **B. cLTP induces biphasic up-regulation of mRNA translation both in soma and dendrites.** Summary of cFRET intensity at baseline, 2, 4, and 6 hours following cLTP activation, as well as 6 hours post-activation after application of GluR1 antagonists 30 minutes prior to activation, in neuronal somata and dendrites. Up-regulation detected at 2 and 6 hours post activation. All comparisons are to t=0h. *** p<0.001, **** p<0.0001 **C. Level of cFRET intensity in somata and dendrites remains unchanged over all time points and following application of GluR antagonists.** Ratio of cFRET intensity in soma and dendrites at baseline, 2, 4, and 6 hours following cLTP activation, as well as 6 hours post-activation after application of GluR antagonists 30 minutes prior to activation.

Interestingly, mRNA translation levels were lower 4 hours after activation compared to the 2 and 6 hour time points. Furthermore, application of DNQX (AMPA receptor antagonist) and APV (NMDA receptor antagonist), together termed GluR antagonists, 30 minutes prior to cLTP activation blocked the up-regulation 6 hours following cLTP (p<0.0001), suggesting that the biphasic up-regulation of protein synthesis is glutamate receptor-dependent, as is known for late-phase LTP (L-LTP, LTP2) (Figure 6B) (67, 106).

In order to compare the results obtained with our new method with an alternative established method, we repeated this experiment using immunocytochemical SUnSET, in which puromycin incorporation into nascent peptides is measured as mean fluorescent intensity normalized to area (Figure 6 – figure supplement 2A). Protein synthesis levels appeared to be elevated at 2, 4, and 6 hours post-cLTP activation (p=0.0029, p=0.0005, and p=0.0006, respectively), with the increase at the 6-hour time point successfully blocked by the application of GluR antagonists 30 minutes before activation (p=0.1415, Figure 6 – figure supplement 2B). These findings bear a resemblance to the results obtained by tRNA-FRET, with the exception of two important points: (1) the 4-hour time point, at which SUnSET experiments reported up-regulation of protein synthesis, obscuring the biphasic pattern observed using tRNA-FRET, and (2) the spatial resolution, and the possibility to conduct this experiment in live cells.

## Discussion

In this study, we employed the tRNA-FRET technique to establish a novel method that enables the visualization and quantification of both tRNA dynamics and mRNA translation at a subcellular resolution (8, 9). We focused on global mRNA translation, whose delicate balance may distinguish between health and disease (10, 11). tRNA-FRET offers specific focus on neuronal sub-compartments and the identification of local translation events with precise temporal and spatial resolution. This capability is especially advantageous in neurobiological research, where the localization of mRNA translation within a single cell and synapse is crucial for proper function (1). Importantly, tRNA-FRET is also suited for live-cell imaging.

The most-studied, unchallenged function of tRNA is its role in mRNA translation. However, tRNA has many cellular functions beyond translation, both regulatory and sensory (6, 7, 15, 107). Nevertheless, tRNA itself has not been studied quite as extensively as many of its cellular targets and regulators, least of all in neurons. Here, we directed our attention to tRNA, viewing its sub-cellular distribution, monitoring its motility in neurons as well as in glia cells, and observing its dynamics.

Through the advantages of labeled tRNA and the tRNA-FRET method, we were able to monitor multiple aspects of neuronal tRNAs and mRNA translation. One of these properties is the sub-cellular distribution of tRNA and its co-localization with translation-related organelles (rough ER) and translation machinery components. By assessing the degree of co-localization of labeled tRNA with the rER in neurons and comparing it with the degree of co-localization in glial cells, we show that the fraction of tRNA co-localized with rER is much lower in neurons compared to glia cells (Figure 1D-E). This finding falls in line with the accepted pattern of cytoplasmic ribosome localization in neurons, where the fraction of ribosomes attached to rER membranes is lower and the fraction of free ribosomes is higher (4) compared to the state in glial cells and other non-neuronal cells (5).

We further examined the nanoscale organization of representative components of translational machinery, namely, tRNA, rpS6, and rpL10a. We were thus able to differentiate between the clustered and diffused fractions of both ribosomal proteins. Through the use of the translation inhibitor puromycin, we determined that the diffused fractions of both ribosomal proteins are the ones engaged in active mRNA translation, as they were both responsive to puromycin as well as co-localized to large tRNA puncta. Importantly, classification of tRNA puncta based on size was shown recently (8), pointing to a translational role specific to the large tRNA puncta. This finding is also consistent with observations that the majority of active ribosomes in neurons are found in the form of free polysomes, unattached to membranes and cellular organelles (45).

Monitoring tRNA mobility in live cortical culture, we observed similar mean instantaneous velocities in both neuronal somata and glial cells, whereas instantaneous velocity was found to be significantly slower in neuronal processes. This could be attributed to the differences between the distinct compartments of neurons, which differ in their cytoskeletal composition (12) and thus may plausibly differ in their modes of transport. Though significantly different from each other, the mean instantaneous velocities we detected in dendrites and in neuronal somata and glial cells correspond with the definition of slow transport (14), which has historically been proposed as the mode of transport for cellular RNA (13). The back-and-forth movement of tRNA puncta along the dendritic shaft observed in our work (Figure 3) highlights the possibility of facilitated transport as opposed to random diffusion. Several mechanisms have been proposed to explain slow transport, one of which claims that molecules tend to randomly switch between anterograde and retrograde transport based on the ratio of kinesin to dynein molecules, thus prolonging the duration of transport (12, 13, 108). These observations pose tRNA-FRET as a promising method for studying tRNA dynamics in neurons in response to varying environmental and cellular conditions.

We briefly examined several aspects of age-dependent differential organization of tRNA in neurons based on numerous previously described findings highlighting the substantial differences between young (immature) and mature neurons, including expression profiles, functionality, and cell durability (21–24). Our data show that tRNA is preferentially localized to the soma in mature neurons (14 DIV) and forms a larger number of aggregates at dendritic branching points, putative “hot spots” of mRNA translation (103) compared to young neurons (7 DIV; Figure 1 – figure supplement 2). These differences may hint at changes in mRNA translation patterns and levels, as well as global regulation of protein synthesis, during neuronal development.

Accordingly, when measuring protein synthesis using tRNA-FRET and comparing the levels of global mRNA translation in young and mature cultured mouse cortical neurons using both tRNA-FRET and SUnSET, we observed a significantly higher level of global translation in young (7 DIV) cultured neurons, in accordance with previous studies (26). We then measured the patterns of neuronal mRNA translation over time following stimulation.

Chemical activation of LTP *in vitro* revealed a pulsatile pattern of global mRNA translation up-regulation that persisted at least six hours post activation. This observation is supported by numerous findings from both *in vitro* (76) and *in vivo* studies (69–75), as well as by the observation of biphasic activation of mRNA translation regulator mTOR (68), that suggest dependency of neuronal processes such as late-phase LTP (L-LTP) and long term memory consolidation upon two or more waves of *de novo* protein synthesis. Conceivably, our results may also be supported by a similar pattern of up-regulation of transcription of Arc mRNA, an immediate early gene (IEG) known to be activated in response to LTP (42, 43), which is synthesized in waves approximately two hours apart and transported into dendrites (25). Notably, this up-regulation of mRNA translation was blocked 6 hours post activation by prior application of GluR antagonists, consistent with reports of GluR-dependent LTP (65–67). These results were further corroborated by immunocytochemical SUnSET.

Examining sub-cellular compartments through the unique benefits of tRNA-FRET, we determined that somatic and dendritic protein synthesis equally contributed to the pulsatile pattern of mRNA translation up-regulation, consistent with previous observations (110), and further strengthened by our presentation of the constant ratio of ∼1 between somatic and dendritic mRNA translation levels regardless of cLTP induction and GluR inhibition.

Research regarding tRNA itself, which has been limited for a long time, has recently attracted renewed attention. Aside from its canonical role as the adaptor that facilitates mRNA translation to proteins, tRNA participates in many additional cellular pathways, including N-terminal conjugation of amino acids to proteins, signal transduction pathways that respond to nutrient deprivation, and regulation of apoptosis (16, 19, 27). As different cell types and cells in different states have different requirements derived from their function and environment, it is highly likely that these cells also differ in their tRNA levels and repertoire, as previously shown (17). Neurons largely differ in these aspects from other cell types. They have been shown to have higher overall expression levels of all tRNAs than almost any other tissue (17), as well as to be enriched with specific tRNA species compared to others, producing a unique tRNA expression profile (17, 18). Moreover, impaired regulation of tRNA biosynthesis and turnover play a direct role in pathology and result in highly complicated clinical phenotypes, including neurodegeneration (18, 20, 28).

Although tRNA is the primary substrate of protein synthesis, mRNA translation is a complex process, and as such, it is possible that other relevant elements may be defective and thus underlie pathology. Disruptions in global mRNA translation stemming from various causes of dysregulation by transcriptional and translational factors are known to be involved disorders such as Fragile X syndrome (33), Rett syndrome (31, 32), Parkinson’s disease (29), Alzheimer’s disease (59), and many more (30). Taken together, these disorders reflect the importance of proper function and proteostasis in neurons. Thus, it is not surprising that a wide variety of methods have been developed in an attempt to precisely monitor and map mRNA translation. These range from early approaches such as labeling with non-canonical amino acids (ncAAs) containing radioisotopes to more contemporary techniques including the ncAA/AHA-based BONCAT (38) and FUNCAT (39), the reporter-mRNA based TRICK (36), the puromycin-based SUnSET (2, 3), the sequencing-based polysome- and ribosome-profiling (37), the Flag tag-based nascent chain tracking (NCT) (35), affinity purification-based TRAP (34), FRET-based smFRET (41), and SunTag-based SINAPS (40, 52).

Each of these methods has its unique advantages, but also its shortcomings, be they the lack of sub-cellular resolution, toxicity, single protein specificity, or inapplicability in live-cell experiments. We found that the innovative tRNA-FRET technique provides information about global mRNA translation at a single-event resolution and is applicable to live cells. An obvious drawback of the method is that the tRNA introduced to the cells is exogenous, and as such is limited in the quantity that can be safely added. Exogenous tRNA constitutes 5-10% of the native cellular tRNA repertoire (44), and thus presumably allows visualizing approximately 5% or less of translation events taking place at any given moment. This concentration is sufficient to detect a signal and to indicate fluctuations in number and intensity in response to stimulation, while simultaneously being harmless to the cells. Moreover, as this proportion remains more or less constant throughout all of our experiments, they are all intercomparable. In the future, we plan to overcome this issue by constructing a viral vector, which will enable monitoring endogenous tRNA.

In summary, neurons transfected with labeled tRNA maintain their functionality and intrinsic properties, enabling monitoring tRNA dynamics in live neurons and differentiating between sub-cellular compartments. Our findings shed light on the dynamics of tRNA molecules in neurons, both spatially and temporally, and provide fine tuning of a research method promising the ability to utilize the precise measurement of mRNA translation events as a tool in the study of translation regulation, regulation through localization and compartmentalization, and potentially in the study of neurodevelopmental and neurodegenerative diseases. Reliable monitoring of protein synthesis in fixed and live cells will enable the characterization of translation profiles of neurons in different states, within sub-cellular localizations such as the synapse, and in response to different stimuli.

The application of the tRNA-FRET method in neuronal primary cultures harbors great potential for the study of neurodevelopmental and neurodegenerative diseases, and it can also be applied to rodent models of disease or to human cells, including stem cells and patient-derived neuronal cells, for patient- and mutation-specific studies.

## Data availability

The link would be provided immediately following initial acception.

## Supplementary data

1. Figure Supplements.

2. Supplementary information.

Supplementary data are available at NAR online.

## Acknowledgements

Arc:dVenus mice were given by Steven Kushner (Department of Psychiatry, Erasmus University Medical Center, Rotterdam, The Netherlands). The authors wish to thank Dr. Iris Alroy (Anima Biotech), Prof. Marcelo Ehrlich (Tel Aviv University), and Dr. Shunit Gal-Ben-Ari (laboratory of K.R.) for critically reading the manuscript.

---

**Figure 1 – figure supplement 1.**
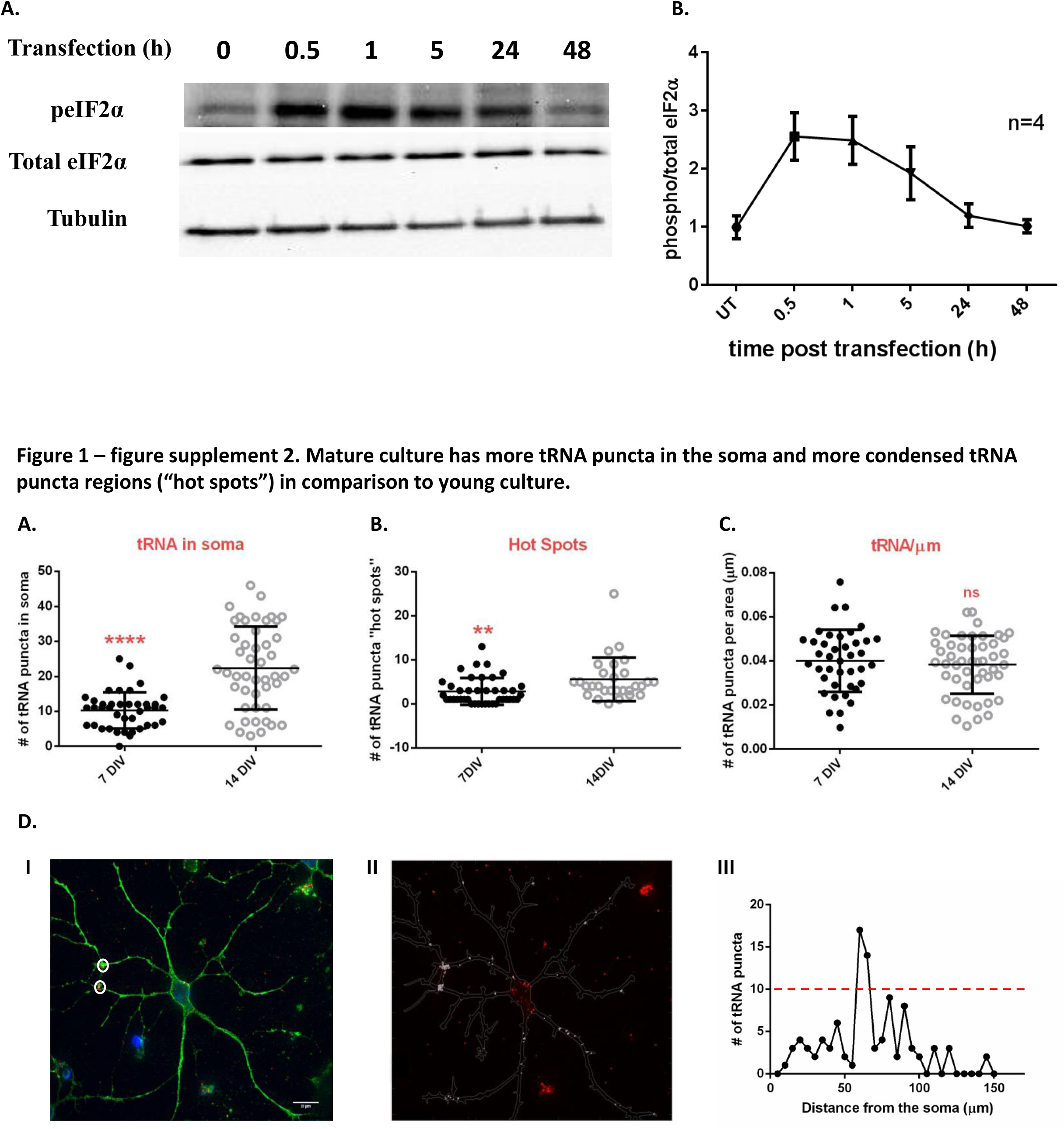
Levels of peIF2α return to baseline 48h following application of tRNA. **A.** Blots of whole cell lysates at different time points (0-48h) following transfection with fluorescently-labeled tRNA. Levels of peIF2α protein were elevated immediately following transfection, and returned to basal conditions after 48h. Total levels of eIF2α protein remained unchanged. Tubulin served as loading control. **B.** Quantification of peIF2α blots shown in A normalized to the total protein levels. n=4.

**Figure 1 – figure supplement 2.**
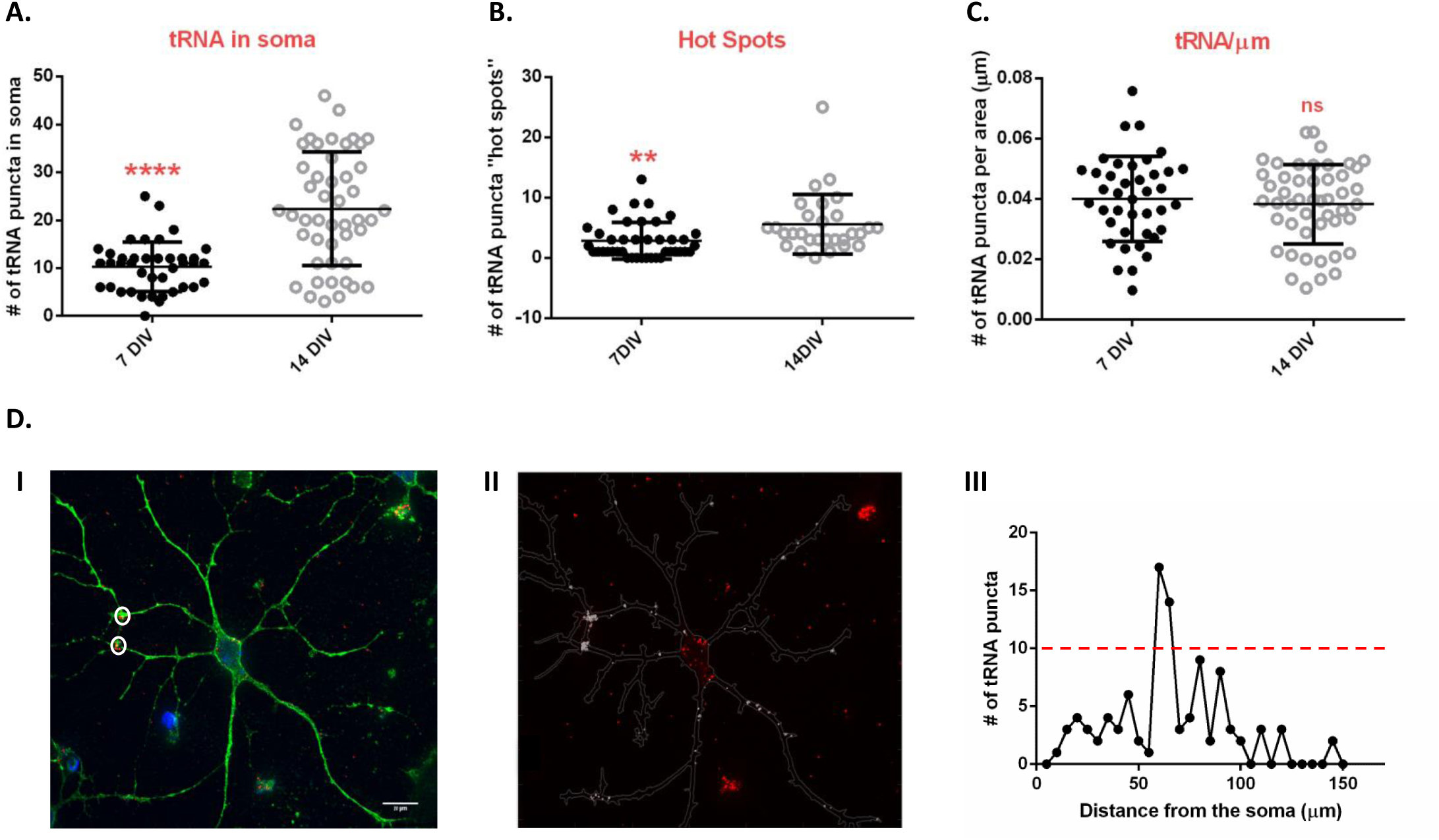
Mature culture has more tRNA puncta in the soma and more condensed tRNA puncta regions (“hot spots”) in comparison to young culture. **A.** Mature culture (14 DIV) has more tRNA puncta in the soma compared to young (7DIV) culture. **B.** Mature culture has more condensed tRNA punctum regions (“hot spots”) compared to young culture. **C.** Normalization of tRNA puncta to dendrite length showed no difference between the two cultures. n _neuron_ ≥30 of 3 independent biological repeats; * p<0.05; **** p<0.0001 **D.** (I) Representative image of a neuron transfected with Cy3-tRNA and stained with Map2-488 and DAPI. Locations of large tRNA puncta (“hot spots”) are circled. (II) Distribution analysis of the image in (I) performed using MATLAB based algorithm, SynD. The outline of the neuron is drawn according to Map2 mask, and tRNA puncta within the neuron are shown in gray. (III) Number of tRNA puncta observed are plotted as a function of distance from the soma. Accumulations of 10 or more tRNA puncta within 5 µm dendrite length were designated “hot spots”.

**Figure 2 – figure supplement 1.**
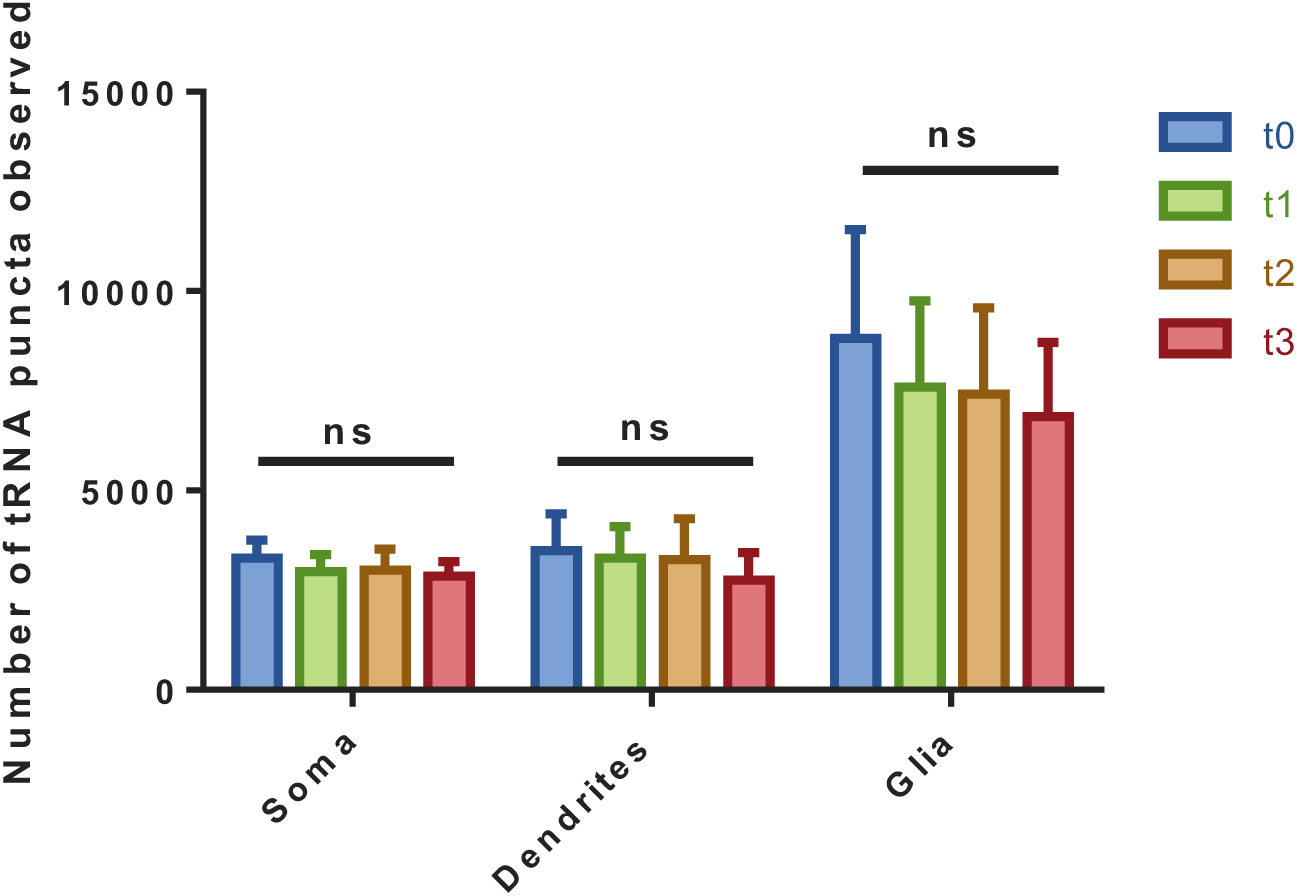
Total amount of tRNA puncta remains unchanged in all measured time points. The total number of puncta observed at all measured time points in somata, dendrites, and glial cells remained unchanged throughout all 4 time points (t0=0-30 min, t1=30-60 min, t2=60-90 min, t3=90-120 min. Puromycin added at t1 and washed out at t2).

**Figure 4 – figure supplement 1.**
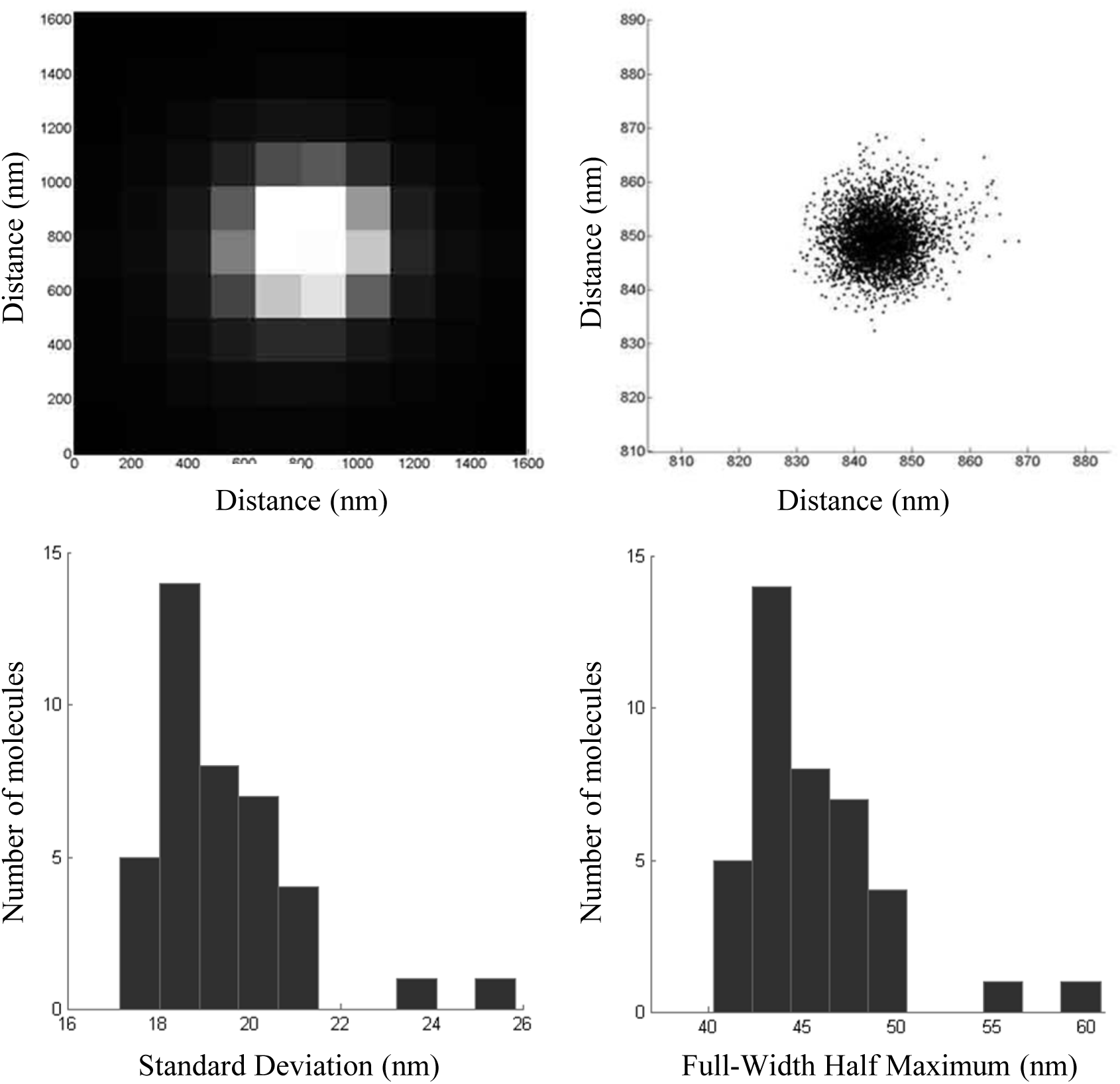
Determination of the spatial resolution of dSTORM. **A.** Point-spread function of a single fluorophore. **B.** Localization pattern of a fluorescent bead that was localized 4000 times. **C.** Histogram of the standard deviation of localizations of 40 single-molecule point-spread function (average standard deviation 19.41 nm). **D.** Histogram of the full-width half maximum (FWHM) of 60 single-molecule point-spread functions (average FWHM 45.6 nm).

**Figure 5 – figure supplement 1.**
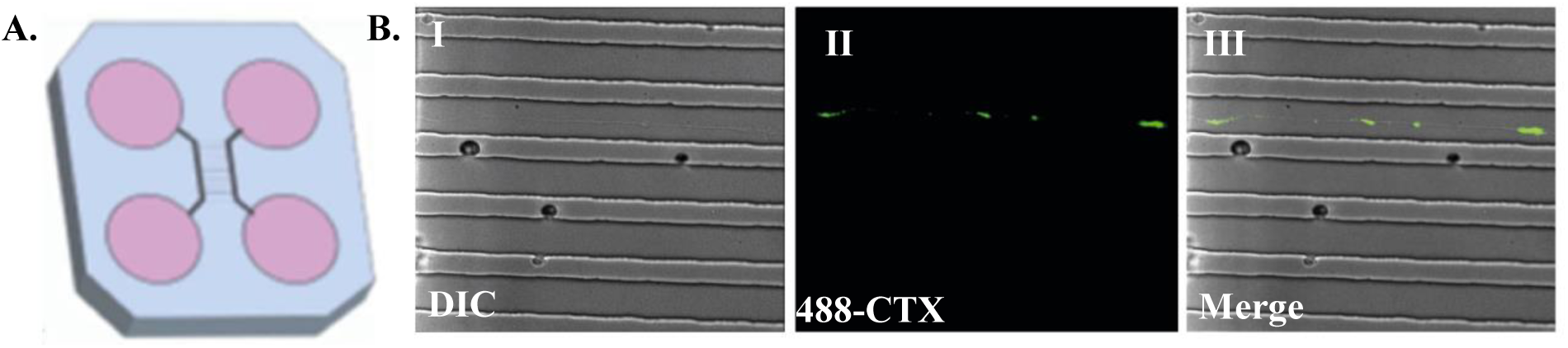
Illustration of a microfluidic chamber. A. Diagram of the microfluidic chamber. B. Representative image of an axon traversing through grooves in the microfluidic chamber in DIC (I), 488-CTX (II), and merged (III).

**Figure 6 – figure supplement 1.**
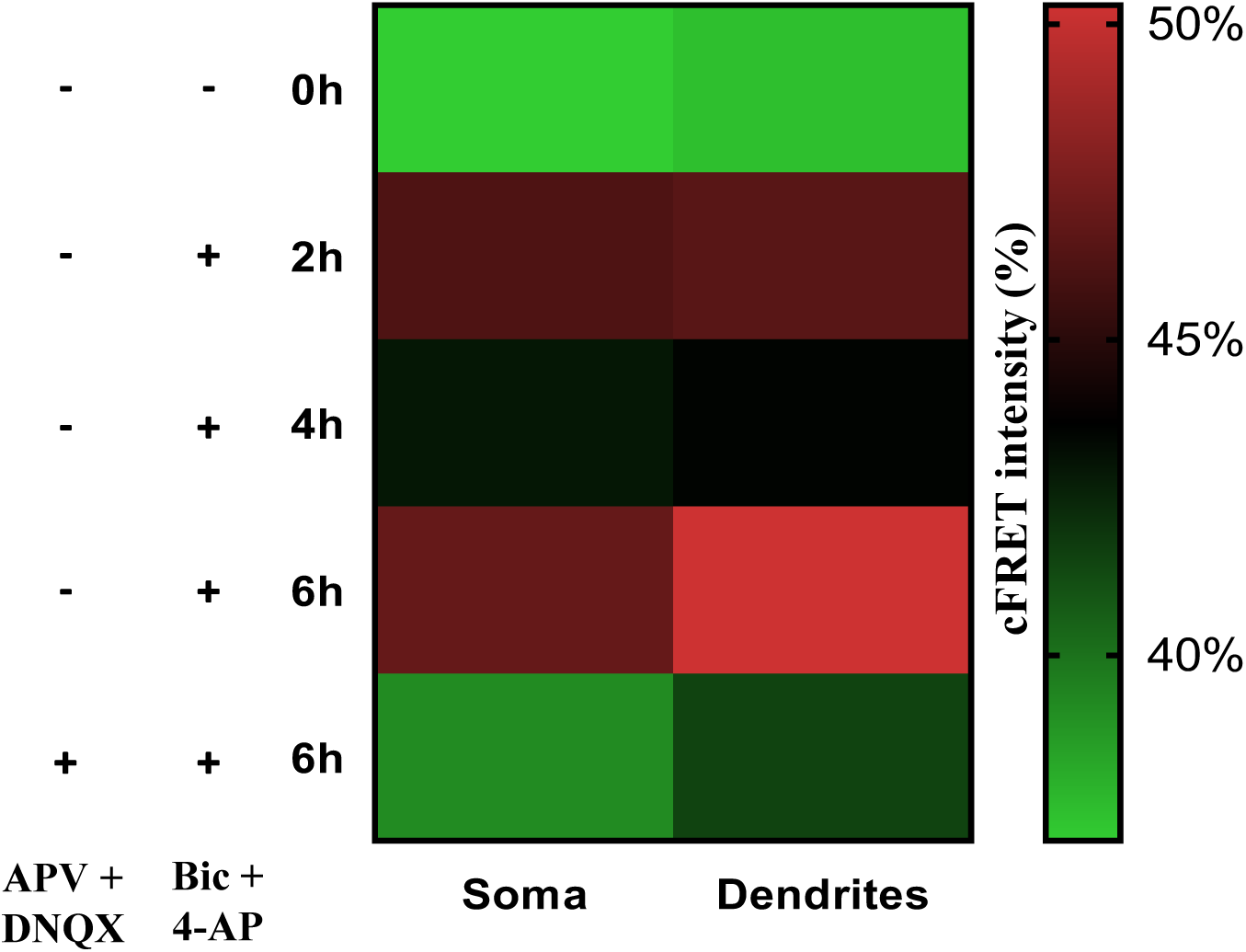
Biphasic pattern of mRNA translation up-regulation is apparent in both neuronal somata and dendrites separately. Mean cFRET intensities in neuronal somata and dendrites at baseline, 2, 4, and 6 hours following cLTP activation, as well as 6 hours post activation after application of GluR1 antagonists 30 minutes prior to activation.

**Figure 6 – figure supplement 2.**
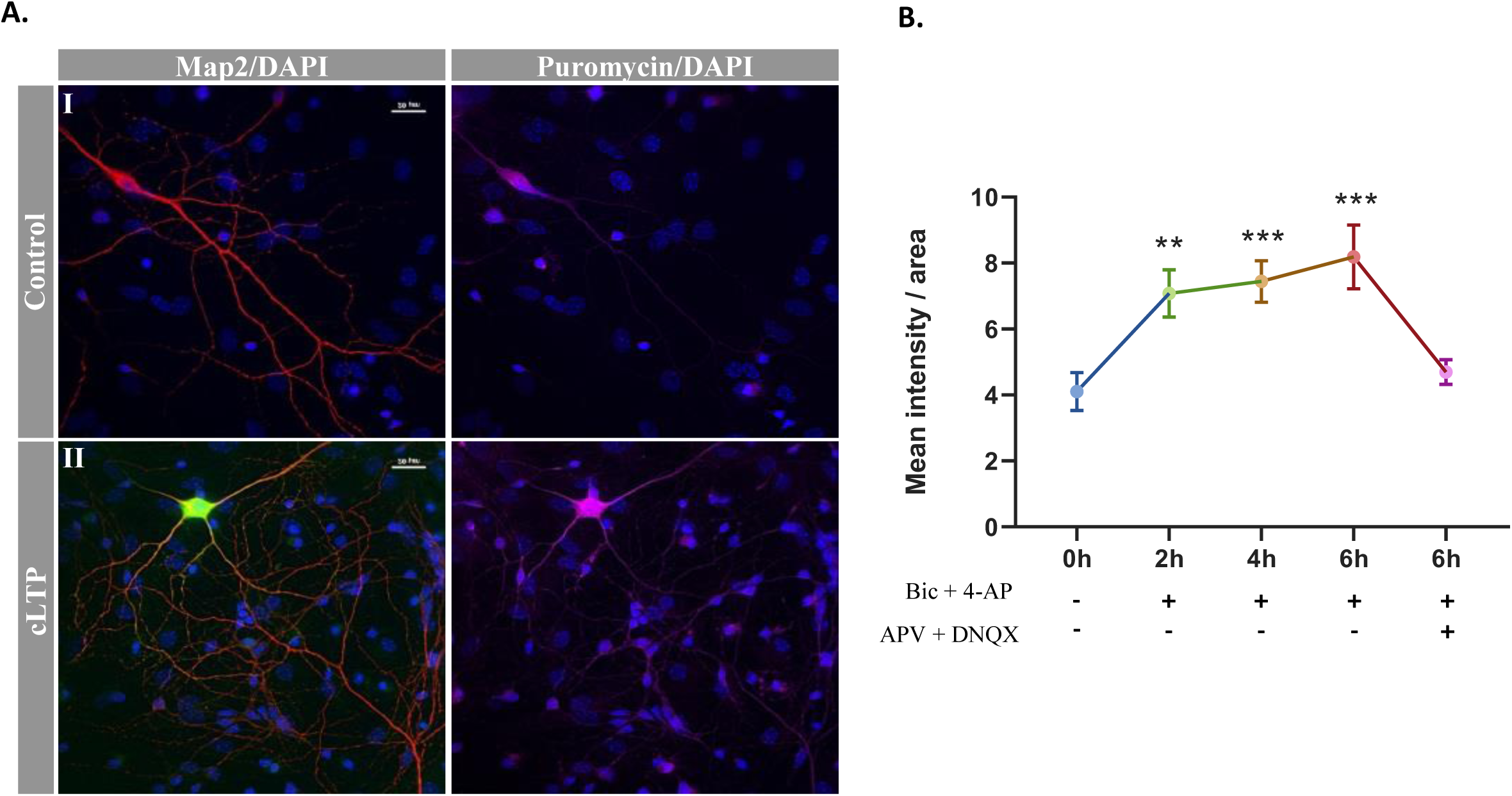
Immunocytochemical SUnSET shows mRNA translation up-regulation following cLTP. **A. cLTP induces time-dependent mRNA translation up-regulation as measured by immunocytochemical SUnSET.** Representative images of Map2-positive neurons at baseline (I) and 6 hours post-cLTP activation (II) stained with αPuromycin antibody. **B. Quantification of protein synthesis levels as measured by SUnSET.** Mean levels of puromycin intensity were normalized to area at baseline, 2, 4, and 6 hours following cLTP activation, as well as 6 hours post-activation after application of GluR1 antagonists 30 minutes prior to activation. Activation of Arc-dVenus served as a positive control for cLTP activation. ** p<0.01, *** p<0.001.

**Supplementary Table 1.** Secondary antibodies used in DNA-PAINT imaging.

| Target | Host | Specification and Commercial Source |
| --- | --- | --- |
| Mouse | Donkey | Donkey Anti-Mouse IgG (H+L)<br>(min X Bov, Ck, Gt, GP, Sy Hms, Hrs, Hu, Rb, Rat, Shp Sr Prot)<br>(#715-005-151, Jackson ImmunoResearch) |
| Rabbit | Donkey | Donkey Anti-Rabbit IgG (H+L)<br>(min X Bov, Ck, Gt, GP, Sy Hms, Hrs, Hu, Ms, Rat, Shp Sr Prot)<br>(#711-005-152, Jackson ImmunoResearch) |

**Supplementary Table 2.** List of docking strand (PD) and corresponding imager strand (PI) used in DNA-PAINT-based imaging study.

| Docking Strand | Imager Strand |
| --- | --- |
| PD-1: Ab-TTTTAGGTAAAG | PI-1: CTTTACCTAA-dye |
| PD-2: Ab– TTTCTTCATTAC | PI-2: GTAATGAAGA-dye |

**Supplementary Table 3.** List of DNA-barcoded labelling agents used in DNA-PAINT imaging.

| Docking Strand | Imager Strand |
| --- | --- |
| Donkey-anti-mouse-PD-1 | PI-1-Atto655 dye |
| Donkey-anti-rabbit-PD-2 | PI-2-Atto655 dye |

## Supplementary Information

### Preparation of DNA-antibody conjugates

1. Antibodies were purchased from commercial vendors (see table S1) and used for conjugation with DNA via thiol-maleimide coupling reactions.

2. Azide or any other preservatives were removed, and the antibody was buffer exchanged to phosphate buffered saline (PBS, pH 7.4) using Zeba spin columns (7000 MWCO).

3. The concentration of the antibody was measured and in a typical conjugation experiment.

4. 100 μg antibody in 75 μL PBS was mixed with 10 eq of maleimide–PEG2–NHS ester in 5 μL of DMF (dimethyl formamide). The solution was then incubated at RT for 2 h.

5. Excess maleimide–PEG2–NHS cross-linker was removed from maleimide-activated antibodies using Zeba spin columns (7000 MWCO, eluent: PBS, pH 7.4) pre– equilibrated with PBS, pH 7.4.

6. In parallel, thiol-modified DNA oligos (10 nmol) were reduced using DTT in 0.1 mL H2O for 1 h at room temperature. The reduced DNA oligos were purified using NAP5 column (GE Healthcare, eluent: PBS, pH 7.4).

7. The maleimide-activated antibodies were mixed with the reduced form of their respective DNA oligos (10 eq) in PBS solution. The reaction was allowed to proceed for 12 h at 4°C.

8. DNA-antibody conjugates were purified and concentrated using Amicon Ultra Centrifugal Filter (100 kDa MWCO).

### Preparation of DNA imager conjugates

1. Amine-modified oligonucleotides were acquired from commercial sources (Integrated DNA technologies) and used for the coupling with NHS ester of Atto 655 fluorophore.

2. Amine-modified DNA (10 nmol, 1 mM in water) was taken in microcentrifuge tube.

3. 1 M NaHCO3 solution was added to the tube for a final concentration of 0.1 M NaHCO3 solution of DNA.

4. Atto 655 NHS ester (2.5 eq, 25 nmol stock in DMF) was added to the DNA solution and stirred at room temperature for 12 h.

5. The conjugated product was purified by reversed phase high performance liquid chromatography (HPLC) using TEAA buffer (buffer A: 5% acetonitrile in 95% 0.1 M TEAA, pH 7.0 and buffer B: 50% acetonitrile in 50% 0.1 M TEAA, pH 7.0) after passing through Zeba spin column.

6. Fluorophore-conjugated oligos were characterized by matrix-assisted laser desorption ionization mass spectrometry (MALDI–MS).

