## Supplementary figures for "Measuring mRNA translation in neuronal processes and somata by tRNA-FRET"

**Figure 1– figure supplement 1. Levels of peIF2α return to baseline 48h following application of tRNA**

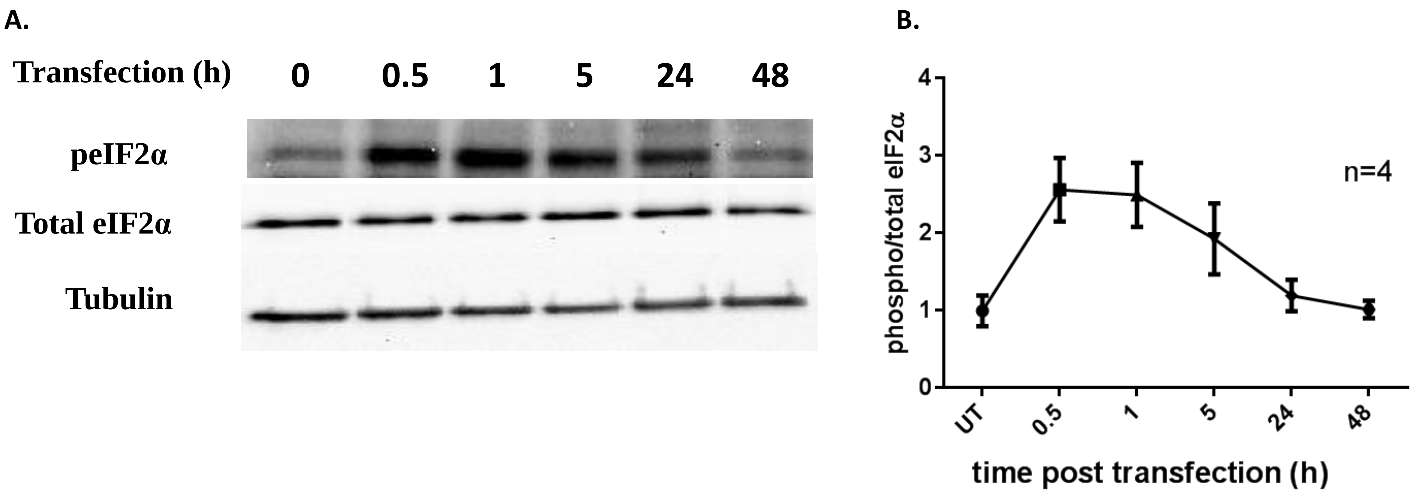


**Figure 1 – figure supplement 2. Mature culture has more tRNA puncta in the soma and more condensed tRNA puncta regions ("hot spots") in comparison to young culture.**

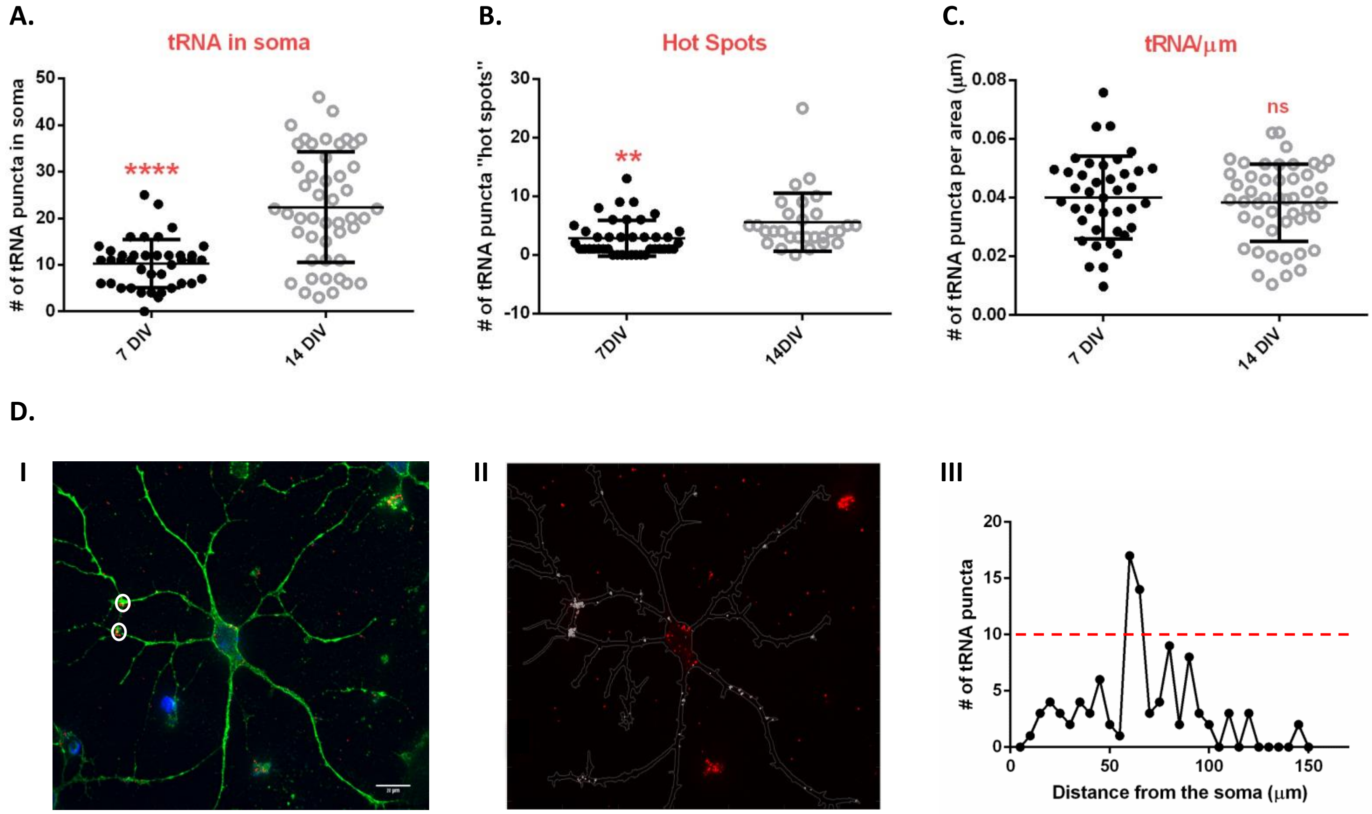

**Figure 2 – figure supplement 1. Total amount of tRNA puncta remains unchanged in all measured time points**

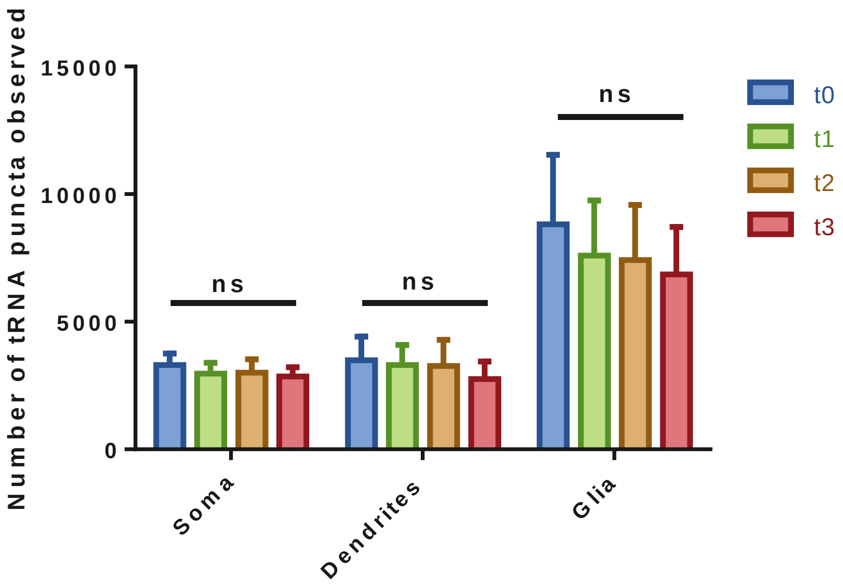

**Figure 4 – figure supplement 1. Determination of the spatial resolution of dSTORM**

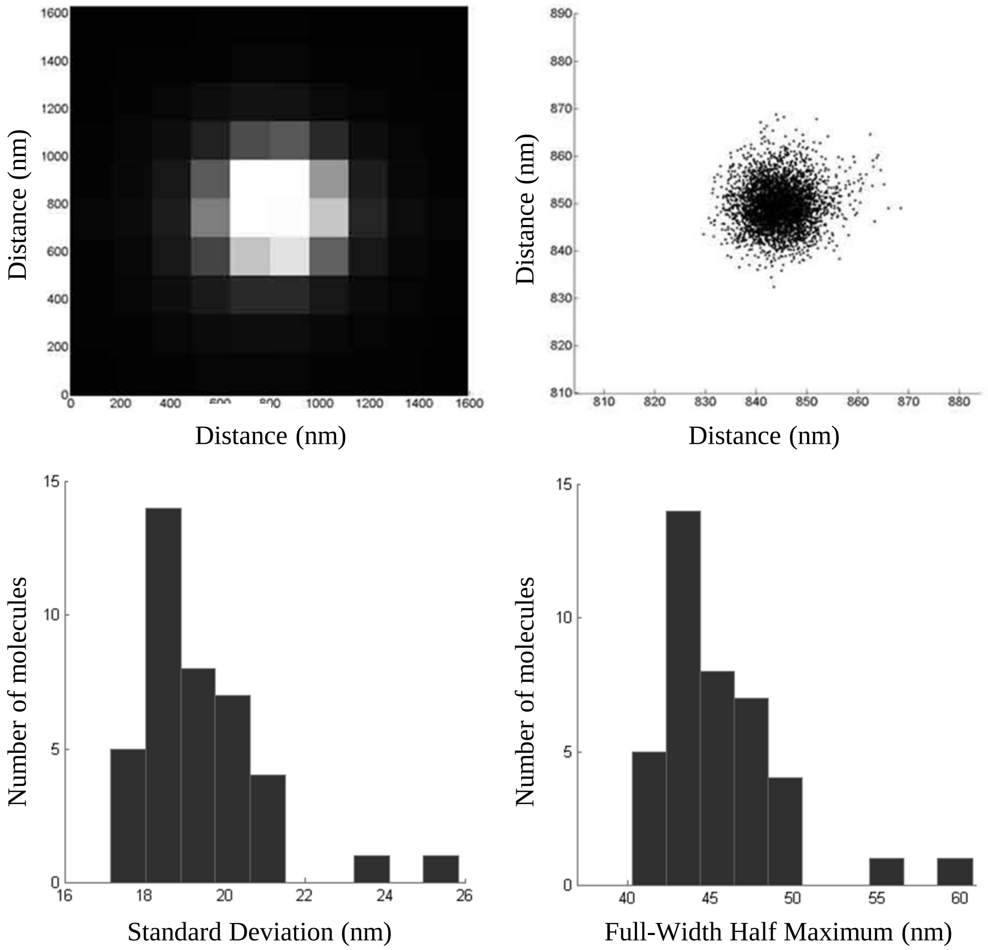

**Figure 5 – figure supplement 1. Illustration of a Microfluidic chamber**

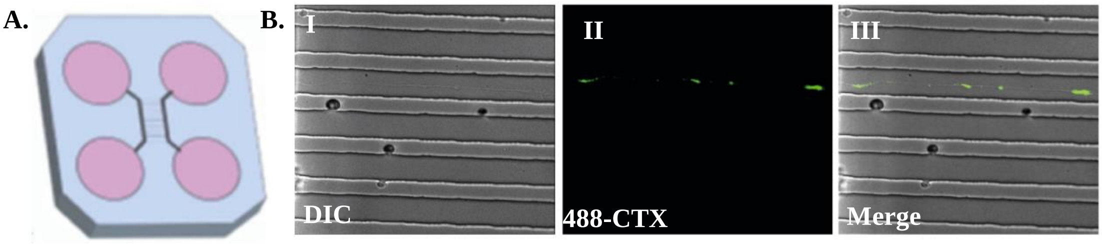

**Figure 6 – figure supplement 1. Biphasic pattern of mRNA translation up-regulation is apparent in both neuronal somata and dendrites separately**

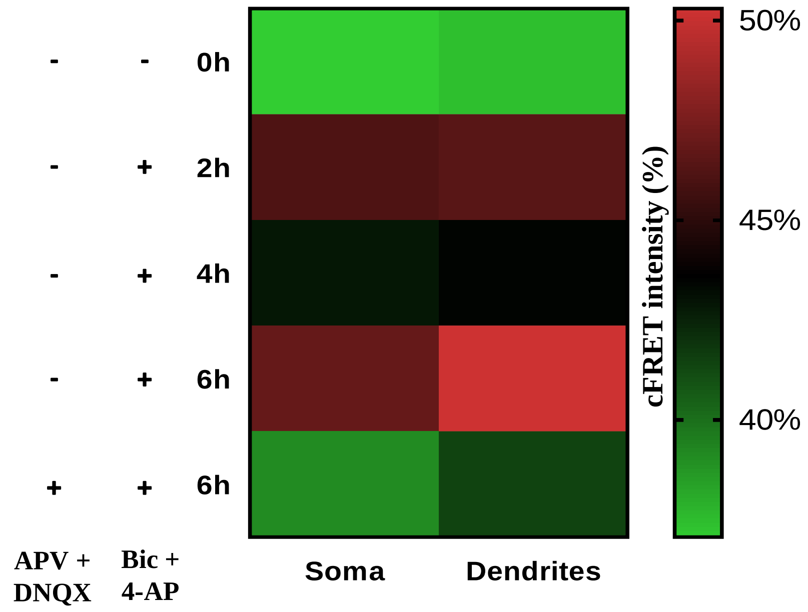


**Figure 6 – figure supplement 2. Immunocytochemical SUnSET shows mRNA translation up-regulation following cLTP**

A.

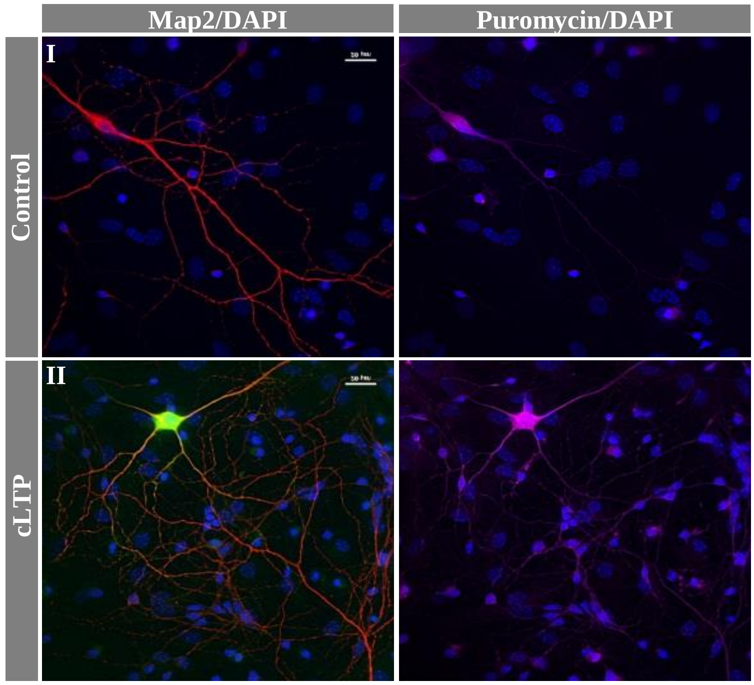


B.

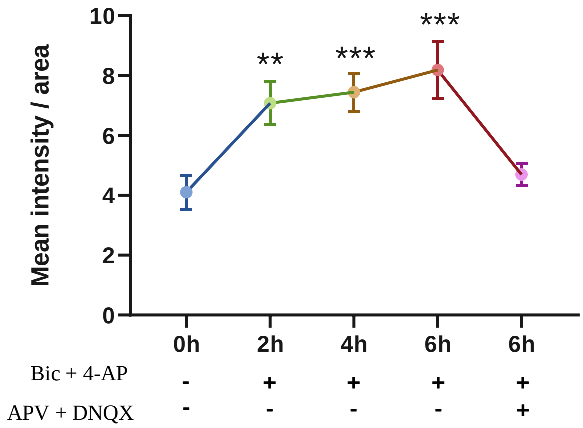
